# Temporal dynamics of CNS cholesterol esters correlate with demyelination and remyelination

**DOI:** 10.1101/2025.09.06.674663

**Authors:** Nishama De Silva Mohotti, Jenna M. Williams, Rashmi Binjawadagi, Hiroko Kobayashi, Disni Dedunupitiya, Jack M. Petersen, Alana Garcia, Meredith D. Hartley

**Affiliations:** Department of Chemistry, University of Kansas, 2030 Becker Drive, Lawrence, Kansas 66047, United States

**Keywords:** Demyelination, remyelination, cholesterol ester, brain, spinal cord, microglia, astrocytes, ACAT1, LCAT, lipid droplets, thyroid hormone agonists

## Abstract

Elevated cholesterol ester levels have been observed in the CNS of patients with neurological diseases, yet the source of cholesterol ester accumulation and whether it is directly linked to demyelination remains undefined. This study investigates the temporal dynamics of cholesterol esters using the *Plp1*-iCKO-*Myrf* mouse model, which features distinct phases of demyelination and remyelination. Our findings reveal that cholesterol ester levels increased with demyelination in both the brain and the spinal cord. In the brain, cholesterol esters declined to normal levels during remyelination, whereas cholesterol esters remained elevated in the spinal cord, which had limited remyelination. Expression of both acetyl-CoA-acyltransferase 1 (ACAT1) and lecithin-cholesterol acyltransferase (LCAT) were elevated during demyelination, suggesting the potential involvement of both proteins in the formation of cholesterol esters. Co-localization studies revealed that ACAT1 is predominantly expressed by microglia and LCAT is predominantly expressed by astrocytes during demyelination highlighting the active roles of glia cells in cholesterol ester metabolism. In addition, we showed that administering the remyelinating drug, Sob-AM2, effectively reduced the level of cholesterol ester accumulation in the brain during demyelination underscoring the potential that manipulating cholesterol ester regulatory pathways may offer for restoring cholesterol homeostasis and promoting remyelination in demyelinating diseases.

## Introduction

Cholesterol is abundantly found in myelin sheaths (∼40% of total lipids) in the central nervous system (CNS) and plays a significant role in maintaining the stability and integrity of myelin membranes, which insulate neurons and facilitate the rapid transmission of nerve impulses.^1–3^ In healthy myelin, cholesterol primarily exists in its unesterified form, while only a minor fraction is present in the esterified form, known as cholesterol esters.^4^ Multiple sclerosis features primary myelin damage caused by an activated immune system, and elevated cholesterol ester levels have been observed in the brain white matter, spinal cord, and cerebrospinal fluid samples collected from people with multiple sclerosis.^5–10^ Increased cholesterol esters have also been detected in the prefrontal cortex of *APOE4* carriers, who have elevated risk of Alzheimer’s disease.^11^ Animal models of multiple sclerosis with demyelination including the cuprizone model^12^ and experimental autoimmune encephalomyelitis (EAE) mouse model have also shown increased levels of cholesterol esters,^13, 14^ and reductions in cholesterol ester levels were observed upon improvement of clinical symptoms.^15^ Collectively, these findings suggest that cholesterol esters accumulate in the CNS as a result of myelin damage and that cholesterol ester hydrolysis may correlate with improved neurological function; however, the cellular and molecular mechanisms governing cholesterol and cholesterol ester metabolism during myelin damage in the CNS have not been well defined.

Cholesterol ester metabolism encompasses synthesis, storage, transportation, and breakdown. Primarily from studies on peripheral tissues, acetyl-CoA-acyltransferase 1 (ACAT1) and lecithin-cholesterol acyltransferase (LCAT) are known to catalyze the formation of cholesterol esters. ACAT1 is a membrane-bound enzyme that is primarily located in the endoplasmic reticulum and regulates intracellular cholesterol homeostasis by catalyzing the esterification of free cholesterol into cholesterol esters, which are stored in the cytoplasm as lipid droplets. Altered ACAT1 expression has been recently reported in neurodegenerative diseases, including Alzheimer’s disease,^16–19^ and blocking ACAT1 activity decreased the production of amyloid plaque formation.^16^ Furthermore, pharmacological inhibition of ACAT1 using ACAT1 inhibitors (CP-113,818 and CI-1011) and genetic ablation of ACAT1 gene have markedly reduced amyloid pathology and cholesterol ester levels in Alzheimer’s disease models.^20–22^

In the periphery, LCAT is primarily synthesized in the liver and plays a crucial role in high-density lipoprotein (HDL) metabolism by converting free cholesterol in circulating lipoproteins into cholesterol esters.^23^ In the CNS, LCAT is predominantly secreted by astrocytes and catalyzes the esterification of cholesterol on apolipoprotein E (APOE) and HDL-like particles.^24^ This action has been linked to a potential role in regulating lipid metabolism in neurodegenerative diseases.^25, 26^

In this study, we utilized a genetically induced demyelination model based on the *Plp1*-iCKO-*Myrf* strain, which features discrete phases of demyelination and remyelination upon induced loss of myelin regulatory factor (*Myrf*) to determine the role of cholesterol esters in demyelination.^27, 28^ In this model, all CNS tissues including brain, spinal cord and optic nerve undergo near complete demyelination followed by robust remyelination in the brain and optic nerve^28, 29^ and chronically impaired remyelination in the spinal cord.^30^ We profiled longitudinal cholesterol ester changes in the brain, spinal cord, and serum in the *Plp1*-iCKO-*Myrf* mouse model. In the brain, cholesterol esters accumulated during demyelination and normalized with remyelination. In contrast, we observed a persistent accumulation of cholesterol esters in the spinal cord even during time points associated with remyelination in the brain. These observations were consistent with the impaired remyelination in the spinal cord previously reported.^30^ Taken together, these results suggest that cholesterol ester accumulation in the CNS correlates with the degree of myelin damage. Furthermore, immunofluorescence analysis revealed that microglial ACAT1 and astrocyte-derived LCAT are both highly expressed during demyelination, suggesting that multiple biochemical pathways may participate in cholesterol ester formation. Finally, treatment with a thyroid hormone agonist, which was previously shown to promote remyelination in this model,^28^ reduced the accumulation of cholesterol esters identifying cholesterol ester metabolism as a potential therapeutic opportunity for promoting remyelination.

## Results and Discussion

### Cholesterol ester levels correlate with the degree of myelin damage

In this study, demyelination in *Plp1*-iCKO-*Myrf* (*Plp1-CreERT; Myrf(fl/fl)*) mice was induced at 8 weeks of age with tamoxifen injections for five consecutive days. Tamoxifen activates Cre recombinase and facilitates cell-specific *Myrf* gene ablation in mature oligodendrocyte cells driven by the *Plp1* promoter.^27, 31^ Loss of the *Myrf* gene leads to oligodendrocyte cell death and progressive loss of myelin, which peaks at ∼8-10 weeks post-tamoxifen. Then, unaffected oligodendrocyte precursor cells (OPCs) proliferate and differentiate to participate in an endogenous remyelination starting at 10 weeks post-tamoxifen.^28^ To represent all stages of active demyelination and remyelination, brain and spinal cord tissues from 6, 12, 18, and 24 weeks post-tamoxifen treatment were used in this study. We also analyzed spinal cord tissues at 30 weeks post-tamoxifen to confirm that limited remyelination in the spinal cord persists at this timepoint.

The time course of demyelination and remyelination of this model has previously been characterized by motor disability (rotarod motor test) and histology (Black-Gold staining).^28, 30^ However, the time course of myelin pathology in the brain and spinal cord was not determined across the full disease course. Representative stained sections from the brain and lumbar spinal cord are presented in Figure 1A and 1C. Threshold analysis was conducted to quantify the percentage of myelin staining for each tissue section. Thresholds were set for each timepoint using *Cre* negative (healthy) tissues to encompass the white matter regions. In the brain, myelin content dropped to 84 % of *Cre* negative levels at week 6 and further to 26 % by week 12, indicating marked demyelination. Partial remyelination was observed at week 18 (52 %) and week 24 (59 %), consistent with myelin repair (Fig. 1A-B). In contrast, the spinal cord exhibited a more severe and persistent loss, with myelin staining declining to 49% of healthy levels at week 6 and to only 1 % by week 30, suggesting widespread demyelination with minimal remyelination (Fig. 1C-D). Interestingly, we observed that the outer regions of the spinal cord white matter tracts (lateral, dorsal, and ventral) had almost no Black-Gold staining, whereas the inner regions had visible staining consistent with either preserved myelin or remyelination; however, all myelin staining levels have greatly reduced intensity compared to healthy spinal cords. These data align with our recent lipidomic study that demonstrated that *Plp1*-iCKO-*Myrf* mice had limited remyelination in the spinal cord compared with the brain.^30^ Overall, these results confirm the robust remyelination in the brain and impaired remyelination in the spinal cord in the *Plp1*-iCKO-*Myrf* model.

**Figure 1.**
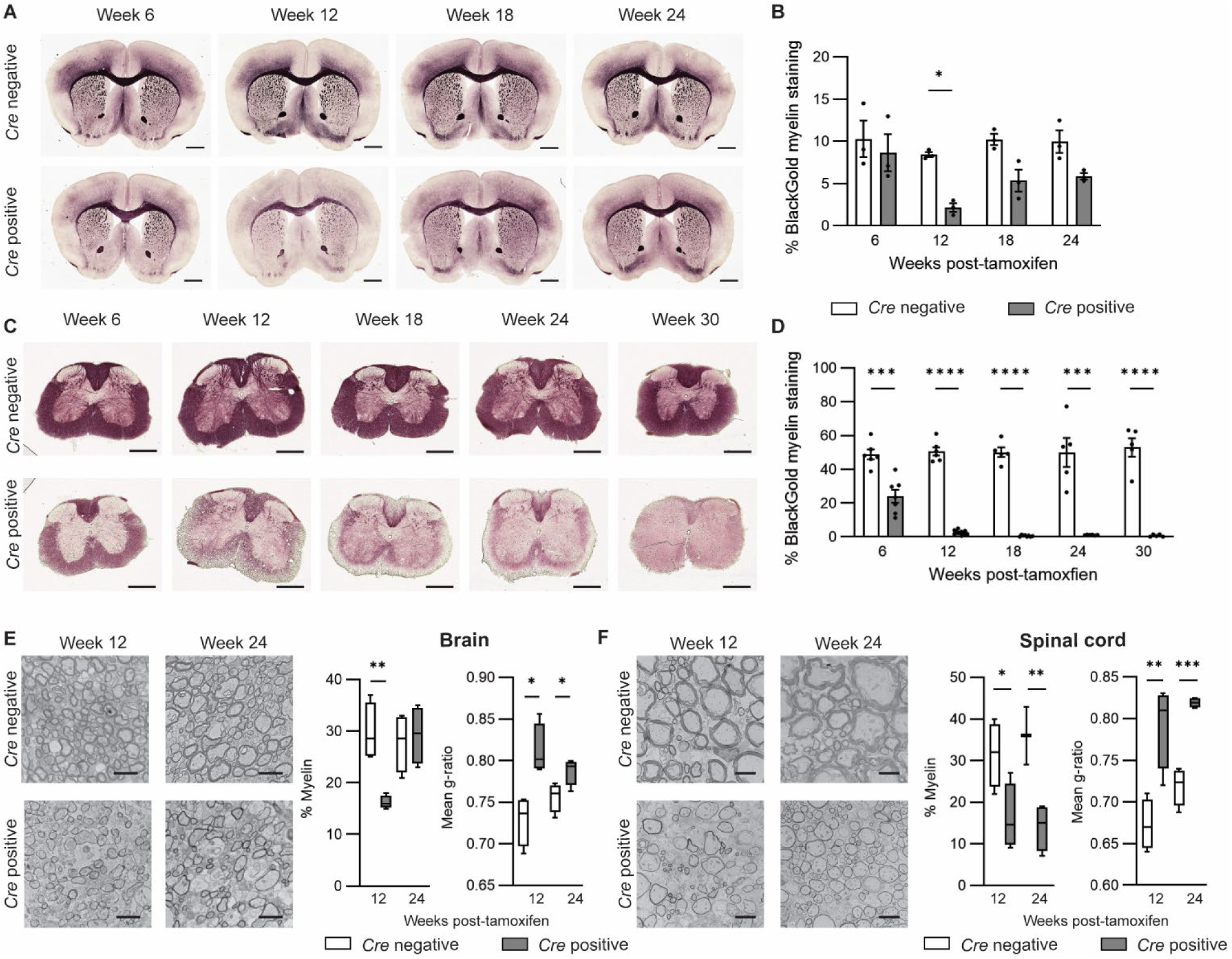
Limited remyelination occurs in the spinal cord relative to the brain. Brain myelin decreases at week 12 and increases by week 24, whereas in the spinal cord, myelin levels are decreased through week 30. (**A** and **C**) Representative Black-Gold-stained tissues from *Cre* negative (healthy) and *Cre* positive (demyelinated) brains and spinal cords from *Plp1*-iCKO-*Myrf* mice. Scale bars in **A** represent 1 mm and in **C** represent 0.5 mm. (**B** and **D**) The staining intensities of the Black-Gold images were quantified by threshold analysis. For the brain, all data represent n = 3 per timepoint. For the spinal cord, the sample numbers are the following: *Cre* negative (week 6 n = 6, week 12 n = 6, week 18 n = 5, week 24 n = 5, and week 30 n = 5) and *Cre* positive (week 6 n = 7, week 12 n = 12, week 18 n = 8, week 24 n = 6, and week 30 n = 5). For all analyses, each sample represents one animal. (**E** and **F**) Representative electron microscopy (EM) images from brain (medial corpus callosum) and lumbar spinal cord at weeks 12 and 24 post-tamoxifen. Scale bars in **E** represent 2 µm and in **F** represent 4 µm. The percentage of myelin by area was determined by threshold analysis. G-ratios were calculated by measuring the axon circumferences and dividing by the outer myelin circumference; at least 90 g-ratios across 3 distinct images (30-50 per image) were quantified for each mouse (n = 3-4 mice for all groups). Statistical analysis was performed with multiple t tests comparing *Cre* negative and *Cre* positive at each time point using a Holm–Šídák correction for multiple comparisons (*P < 0.05, **P < 0.01, ***P < 0.001, ****P < 0.0001).

To quantify remyelination, electron microscopy (EM) analysis was performed on the corpus callosum in the brain and on the lumbar spinal cord. The hallmark of remyelination is thinly remyelinated axons, and “thin” myelin sheaths are observed in both the brain and spinal cord indicating that some remyelination has occurred (Fig. 1 E-F). The percentage of myelin across the EM images was quantified by threshold analysis and was consistent with the BlackGold staining. The brain shows a drop in myelin staining at week 12 followed by an increase to normal levels at week 24. In contrast the spinal cord shows a drop in the myelin staining at week 12 that persists to week 24. In addition, the myelin thickness was quantified using g-ratios (the ratio of the inner axon circumference to the outer myelin circumference). The mean g-ratio is higher in the demyelinated mice at both timepoints and in both tissues. Brains show a much smaller difference in mean g-ratio at week 24 as compared to week 12 consistent with the successful remyelination. In contrast, spinal cord shows a large difference between healthy and demyelinated g-ratios that continues through week 24. The spinal cord features larger axons with thicker myelin, and thus, the extent of myelin loss is more dramatic as compared to the brain when examining either the percentage of myelin or the g-ratios.

To quantify cholesterol ester dynamics during demyelination and remyelination, we used direct infusion triple quadrupole mass spectrometry to measure cholesterol esters in the brain and spinal cord. In the brain, a ∼20-fold increase of total cholesterol ester levels was observed at week 6 during active demyelination that peaked at week 12 with a ∼30-fold increase (Fig. 2A). At week 18, the cholesterol esters decreased reaching near normal levels at week 24, which correlated with the observed robust remyelination (Figure 1A-B). Spinal cord exhibited a larger (∼130-fold) in total cholesterol ester levels (Fig. 2C), peaking at week 18, and remained elevated through week 30 (∼115-fold), which matched the observed persistent myelin loss (Figure 1C-D). In contrast to the CNS, serum cholesterol ester levels did not vary at weeks 6, 12, or 18 post-tamoxifen (Figure S1).

**Figure 2.**
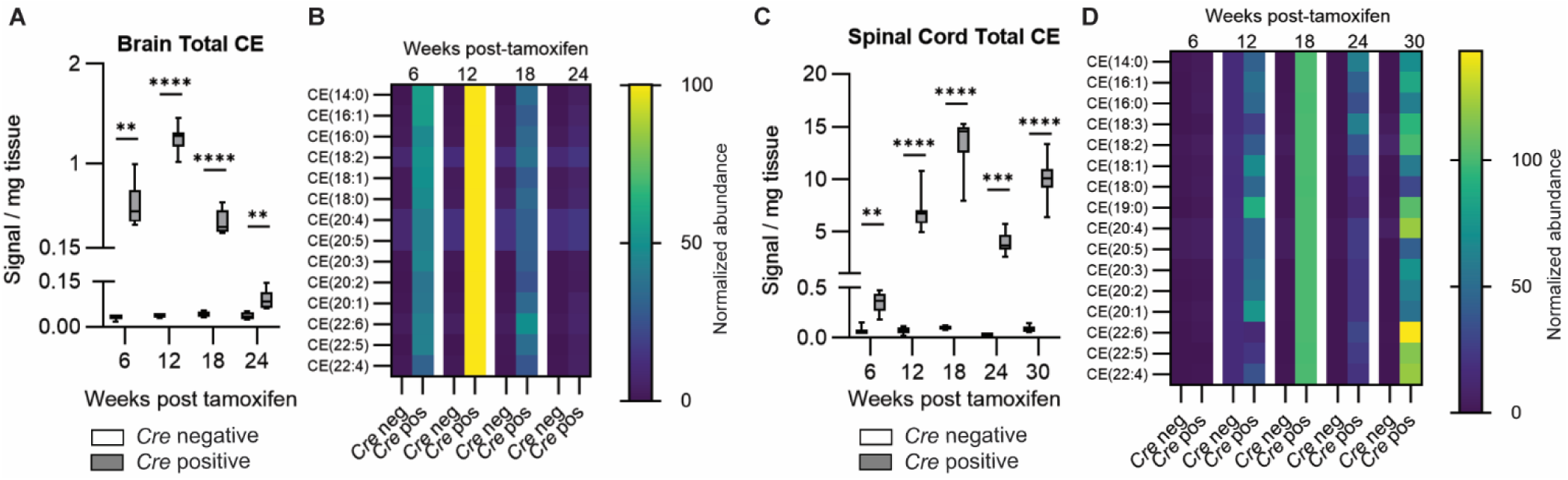
Cholesterol ester levels increase during demyelination and decrease during remyelination. Cholesterol esters were quantified using mass spectrometry. (**A** and **C**) Total cholesterol ester levels for *Cre* negative and *Cre* positive brain (**A**) and spinal cord (**C**) tissues. Individual cholesterol esters were measured and summed to determine the total levels. Box and whisker plots show the mean, minimum and maximum. Statistical analysis was performed with multiple t tests comparing *Cre* negative and *Cre* positive at each time point using a Holm–Šídák correction for multiple comparisons (**P < 0.01; ***P < 0.001; ****P < 0.0001). (**B** and **D**) Heatmaps of the individual cholesterol esters for brain (B) and spinal cord (D). The levels at peak cholesterol ester accumulation (week 12 for brain or week 18 for spinal cord) were set to 100% for each lipid and all other samples were normalized based on the peak level. For **A**-**D**, sample numbers were: brain *Cre* negative, 6 weeks n = 3, 12 weeks n = 5, 18 weeks n = 6, and 24 weeks n = 4; *Cre* positive, 6 weeks n = 8, 12 weeks n = 8, 18 weeks n = 8, and 24 weeks n = 7; spinal cord *Cre* negative, 6 weeks n = 3, 12 weeks n = 5, 18 weeks n = 6, 24 weeks n = 4, and 30 weeks n=7; *Cre* positive, 6 weeks n = 8, 12 weeks n = 8, 18 weeks n = 9, 24 weeks n = 7, and 30 weeks n = 7. Abbreviations: CE, cholesterol esters.

A direct comparison of total cholesterol ester profile (Figure 2A and 2C) to the Black-Gold histological staining (Figure 1A-D) clearly demonstrates that cholesterol ester levels are inversely correlated with degree of myelin damage. The cholesterol ester data coupled with the histology demonstrate that elevated cholesterol esters in the CNS are a marker of demyelination. Furthermore, robust remyelination in the brain is accompanied by the decline in total cholesterol ester levels, suggesting that cholesterol ester accumulation is cleared during myelin repair and may be a critical step for successful remyelination.

We used gas chromatography-mass spectrometry to quantify total cholesterol levels at the time points of maximum cholesterol ester accumulation in the demyelinated *Cre* positive tissues. At week 12 in the brain, cholesterol esters comprise 13 % of the total cholesterol/cholesterol ester population, and at week 18 in the spinal cord, cholesterol esters comprise 24 % of the total cholesterol/cholesterol ester population (Table S1). Both amounts are over 20-fold increases from the normal percentage of cholesterol esters in healthy CNS tissues (< 0.5 %). Thus, there is a substantial perturbation in cholesterol metabolism and storage during myelin damage.

### Polyunsaturated fatty acid cholesterol esters (PUFA-CEs) remain significantly elevated in the spinal cord with impaired remyelination

To visualize how individual cholesterol ester species change during the disease course, heatmaps showing the levels of individual cholesterols esters normalized to the peak accumulation are shown in Figure 2B for brain and Figure 2D for spinal cord. The absolute levels of the different cholesterol ester species reflect the overall abundance of the fatty acids in the CNS and are reported in Figures S2-S3.

In the brain, most cholesterol ester species peaked at week 12, coinciding with peak demyelination followed by a decline in week 18 and week 24 with remyelination. The most abundant cholesterol ester species (Figure S1), such as CE (18:1), CE (16:0), and CE (16:1), substantially increased with demyelination but decreased back to normal levels by week 24 with remyelination. Cholesterol esters derived from polyunsaturated fatty acids (PUFA-CEs) like CE (22:6), CE (20:4), CE (20:5), CE (22:4), CE (20:3), and CE (18:2) also significantly increased with demyelination. PUFA-CEs have been linked to active demyelination and impaired cholesterol processing by microglia and astrocytes.^32^ Therefore, the marked reduction of these PUFA-CEs by week 24 further confirms the robust remyelination and restoration of lipid metabolism in the brain.

In the spinal cord, most cholesterol ester species peak at week 18, aligning with the delayed onset of demyelination in the spinal cord (Figure 2D and S3). Consistent with the observations in the brain, all cholesterol esters species showed a significant increase with demyelination. However, in the spinal cord, most of the monounsaturated and polyunsaturated fatty acid cholesterol esters were still elevated through week 30 and several species (CE (20:4), CE (22:4), CE (22:5), and CE(22:6)) had increased above week 18 levels despite week 30 having overall lower levels of cholesterol esters. The continued elevation of PUFA-CEs is consistent with impaired remyelination and suggests disrupted lipid homeostasis.

Our results suggest that in the *Plp1*-iCKO-*Myrf* mouse model, cholesterol ester levels closely correlate with myelin damage and repair. The brain had dynamic cholesterol ester levels that peaked during demyelination and gradually decreased with remyelination, implying effective cholesterol ester clearance with robust remyelination. In contrast, the spinal cord showed persistent accumulation of cholesterol esters, particularly polyunsaturated cholesterol esters, correlating with persistent myelin damage and impaired remyelination. Taken together, these observations highlight the importance of CNS cholesterol ester formation and breakdown in the CNS response to myelin damage. The lack of remyelination in the spinal cord underscores that the clearance of accumulated cholesterol esters is crucial for successful remyelination.

### Early microglial activation correlates with effective cholesterol ester clearance and robust remyelination

During demyelination, damaged myelin is released into the extracellular matrix where it is phagocytosed by microglia and other macrophages in the CNS.^33^ Previous studies have established the importance of microglial activation, phagocytosis activity, and microglia associated gene expression in clearing myelin debris and promoting remyelination.^34, 35^ Myelin and myelin debris are rich in free cholesterol, and microglia and infiltrating macrophages act as primary phagocytes in the CNS. Several lines of evidence suggest microglia may be involved in the conversion of free cholesterol in myelin to cholesterol esters during demyelination.^2^ ACAT1 is well studied in peripheral macrophages, where it is known to convert cholesterol to the esterified form.^36^ Further, treatment of bone marrow derived macrophages^37^ or microglial cell lines^38^ with myelin increased cholesterol ester synthesis that was blocked by treatment with ACAT1 inhibitors. Inducing demyelination *in vivo* in the cuprizone model of demyelination revealed that cholesterol esters accumulate in microglia,^12^ further highlighting the role that microglia may play in myelin debris processing and cholesterol ester formation during demyelination.

To examine the role of microglia in our model, we performed immunofluorescence for IBA1 to image microglia in the brain and spinal cord tissues across multiple timepoints (Figure 3). In the brain, we examined the corpus callosum, and we observed the most significant accumulation of microglia at week 6 during active demyelination prior to peak demyelination at week 12 (Figure 3A and 3E). This observation suggests that microglia and macrophages have an immediate response to the accumulation of myelin debris which may be independent of the overall extent of myelin damage. It also suggests that early activation of microglia may be critical for initiating successful remyelination. In contrast, in the spinal cord, the maximum accumulation of microglia and macrophages occurred much later at week 18 (Figure 3B and 3G). The delayed activation of microglia in the spinal cord may be insufficient for effective myelin debris clearance which could contribute to the limited myelin repair.

**Figure 3.**
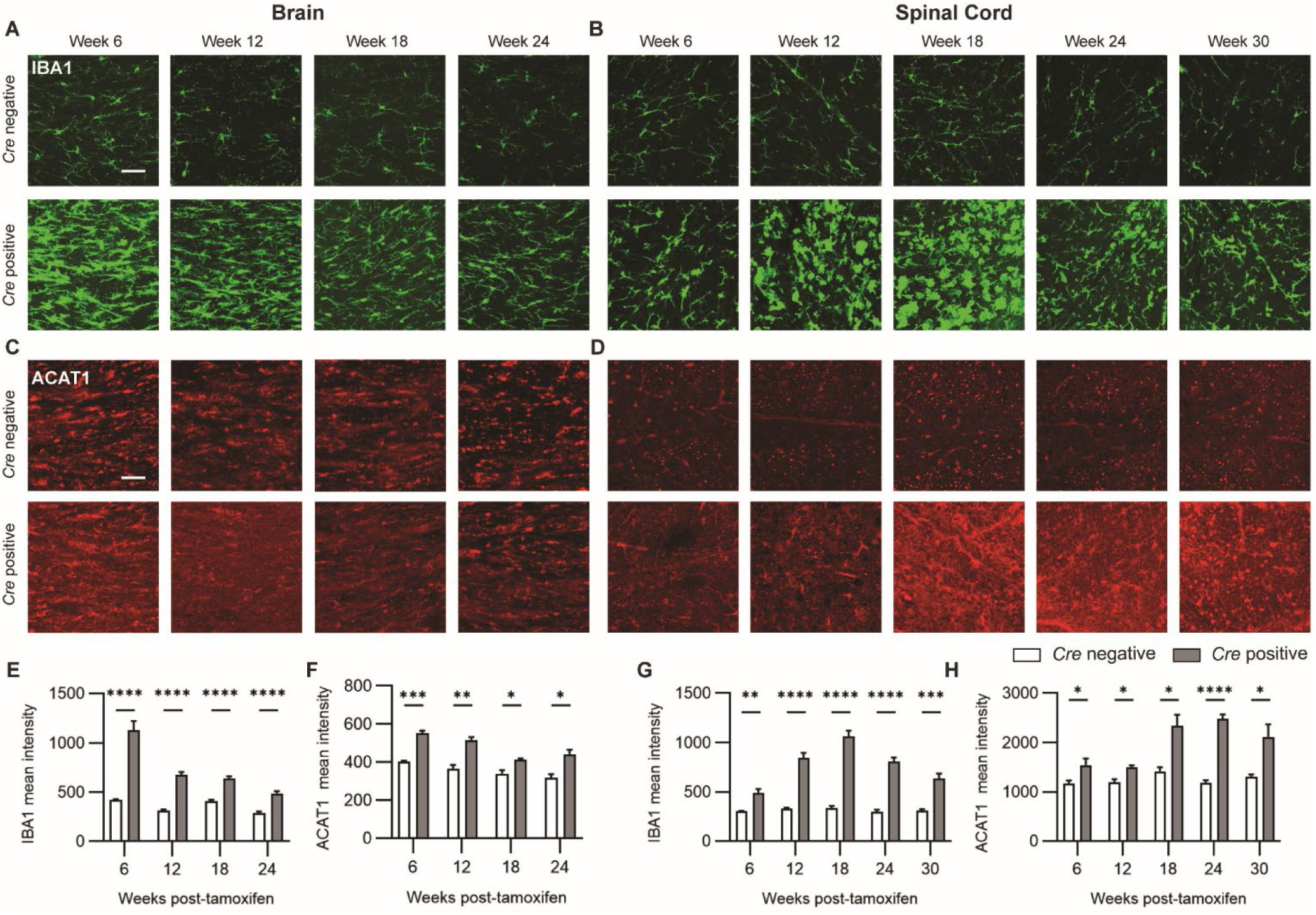
Microglial activation parallels ACAT1 expression in both brain and spinal cord. (**A** and **C**) Representative immunofluorescence images in the brain medial corpus callosum. Brain sections were stained with antibodies for IBA1 (green) and ACAT1 (red). (**E** and **F**) Mean intensities in the corpus callosum are shown. For IBA1 analysis, the number of animals analyzed for each group were the following: *Cre* negative, 6 weeks n = 5, 12 weeks n = 6, 18 weeks n = 6, and 24 weeks n = 5; *Cre* positive, 6 weeks n = 5, 12 weeks n = 6, 18 weeks n = 5, and 24 weeks n = 7. For ACAT1 analysis, n=4 mice were analyzed for all groups. (**B** and **D**) Representative immunofluorescence images in the dorsal region of the lumbar spinal cord. Sections were stained with antibodies for IBA1 (green) and ACAT1 (red). (**G** and **H**) Mean intensities in the dorsal white matter are shown. For IBA1 analysis, two spinal cord sections were stained and the intensities averaged for each animal. The number of animals analyzed were the following: *Cre* negative, 6 weeks n = 5, 12 weeks n = 5, 18 weeks n = 5, 24 weeks n = 4, and 30 weeks n = 5; *Cre* positive, 6 weeks n = 7, 12 weeks n = 8, 18 weeks n = 7, 24 weeks n = 8, and 30 weeks n = 5. For ACAT1 analysis, n = 3-4 mice were analyzed for all groups. Statistical analysis was performed with multiple t tests comparing *Cre* negative and *Cre* positive at each timepoint using a Holm–Šídák correction for multiple comparisons (*P < 0.05; **P < 0.01; ***P < 0.001; ****P < 0.0001). For all images, scale bars indicate 50 µm.

In healthy brain and spinal cord tissues, microglia morphology resembled a ramified, resting state, as expected. However, microglia in demyelinating mice had altered morphologies. Brain microglia appeared partially activated, whereas spinal cord microglia showed an ameboid morphology consistent with a vulnerable and disease-associated microglia state (Figure 3A-B).^33^ A possible explanation of the highly altered microglial state is that spinal cord has more myelin per tissue weight relative to brain, and thus spinal cord microglia are exposed to larger quantities of myelin debris during demyelination.

When myelin debris are phagocytosed by microglia, both excess cholesterol and excess free fatty acids in the engulfed myelin debris can be cytotoxic, and microglia can synthesize neutral cholesterol esters to store in lipid droplets.^12, 39^ The enzymatic pathways responsible for converting cholesterol into cholesterol esters are well-characterized in peripheral tissues and in certain neurodegenerative diseases like Alzheimer’s disease. However, these pathways are not well understood in the context of demyelination. ACAT1 is a transmembrane protein in the endoplasmic reticulum that esterifies cholesterol using acyl-CoA.^38^ We used immunofluorescence to measure how ACAT1 levels align with the fluctuations in cholesterol ester levels (Figure 3C-D). Both brain and spinal cord ACAT1 protein levels correlate with the changes in microglia IBA1 staining (Figure 3E-H), suggesting that microglial ACAT1 likely contributes to cholesterol ester accumulation during demyelination.

Since ACAT1 is not microglia specific and can be expressed by other cell types in the CNS, we performed a co-localization analysis of ACAT1 in microglia. Results revealed that 30% of the brain ACAT1 is expressed in microglia and 54% of the spinal cord ACAT1 is expressed by microglia during demyelination (Figure 4). Although this does not preclude the possibility that ACAT1 in other cell types may also be contributing to cholesterol ester formation, it suggests that microglia ACAT1 is likely a source.

**Figure 4.**
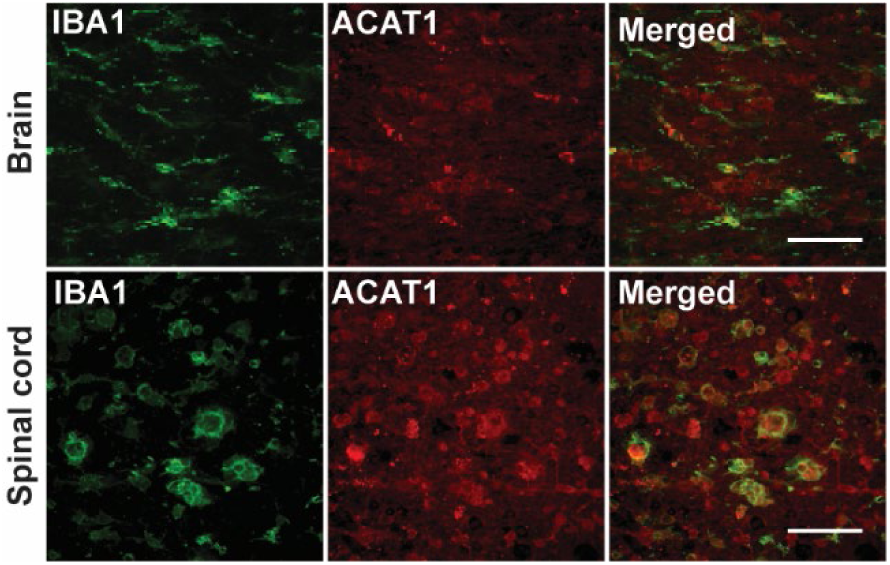
ACAT1 is expressed in microglia during demyelination. ACAT1 expression is co-localized with microglia in brain and spinal cord. Representative brain and spinal cord immunofluorescence images used for co-localization analysis of ACAT1 in microglial cells. Scale bars represent Sections were stained by directly labeling the primary antibodies. ACAT1 was directly conjugated with Mix-n-Stain CF555 (red) and IBA1 with Mix-n-Stain CF647 (green). Threshold analysis was carried out to quantify the percentage of ACAT1 that overlapped with IBA1 staining. Only *Cre* positive mice used for this analysis. Mice numbers: Brain 6 weeks n = 4, spinal cord 18 weeks n = 4.

To further explore the role of microglia in cholesterol ester formation, we stained for neutral lipid droplets in the brain and spinal cord tissues using Lipidspot 610 and Lipidspot 488 respectively (Figure 5A-B). Compared to *Cre* negative samples, *Cre* positive samples showed an accumulation of lipid droplets both in the brain and in the spinal cord during demyelination, with the spinal cord showing larger increases in lipid droplets (Figure 5C-D). Co-staining lipid droplets with microglia and astrocytes revealed that lipid droplets are specifically located in microglia (Figure 5E). The presence of lipid droplets in microglia provides support for an ACAT1-based mechanism where cholesterol esters are formed in microglia during demyelination and are stored in lipid droplets. Our observations align with a previous report showing that inducing demyelination with lysolecithin in ACAT1-deficient mice reduces the formation of lipid droplet– containing microglia and impairs remyelination.^37^

**Figure 5.**
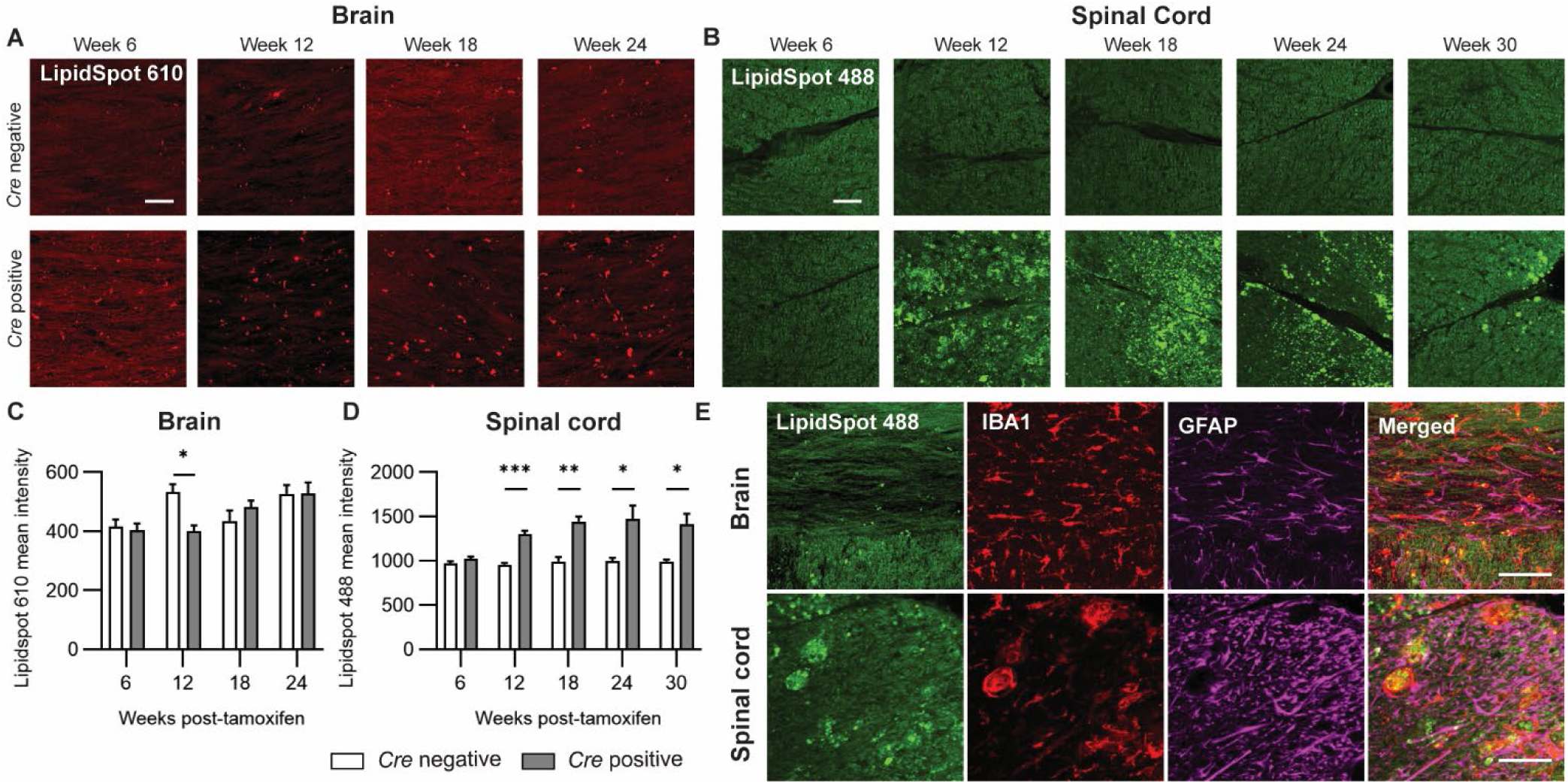
Lipid droplets form in microglia during demyelination. Lipid droplets increase during demyelination in the brain and spinal cord. (**A** and **B**) Neutral lipid droplets were stained using LipidSpot 610 in the medial corpus callosum of the brain and LipidSpot 488 in the ventral spinal cord. (**C** and **D**) A threshold analysis was carried out to quantify the mean intensity of the lipid droplets (n = 4 for all groups). Statistical analysis was performed with multiple t tests comparing *Cre* negative and *Cre* positive at each time point using a Holm–Šídák correction for multiple comparisons (*P < 0.05; **P < 0.01; ***P < 0.001). (**E**) Co-staining shows that lipid droplets accumulated in microglia and not astrocytes in the brain and spinal cord. Fluorescence stain LipidSpot 488 was used to stain lipid droplets (green) with antibodies for IBA1 (microglia, red) and GFAP (astrocytes, magenta). For all images, scale bars indicate 50 µm.

We also observed that lipid droplet accumulation in the spinal cord localized to the outer portions of the ventral and dorsal white matter (Figure S4). In the Black-Gold myelin analysis, we observed that the outer regions of the ventral, dorsal and lateral white matter had little myelin staining relative to the inner white matter regions (Figure 1). Thus, the regions of lipid droplet accumulation in the spinal cord overlapped with the regions of maximum demyelination (Figure S4) suggesting a strong correlation between the lipid droplet accumulation and demyelination. One possible explanation is that microglia in the outer white matter of the spinal cord may have a different phenotype and are unable to clear accumulated cholesterol esters resulting in remyelination failure, but more investigation is needed to understand this observation.

### Overexpression of LCAT suggests a role for astrocytes in cholesterol ester formation

Next, we investigated the cellular response of astrocytes during demyelination and remyelination (Figure 6) using GFAP as a marker for immunofluorescence studies. Astrocyte accumulation and expression of GFAP peaks at week 12 in the brain and reduces during remyelination (Figure 6A and 6E). The spinal cord showed astrocyte accumulation that persisted through week 30 (Figure 6B and 6G). Astrocytes are also the primary source of a second enzyme that can form cholesterol esters known as lecithin-cholesterol acyltransferase (LCAT).^24^ LCAT is primarily synthesized in the liver and acts directly on cholesterol in apolipoprotein particles using phosphatidylcholine (lecithin) as a fatty acid source. LCAT is one of the major cholesterol esterifying enzymes in peripheral tissues maintaining cholesterol homeostasis.

**Figure 6.**
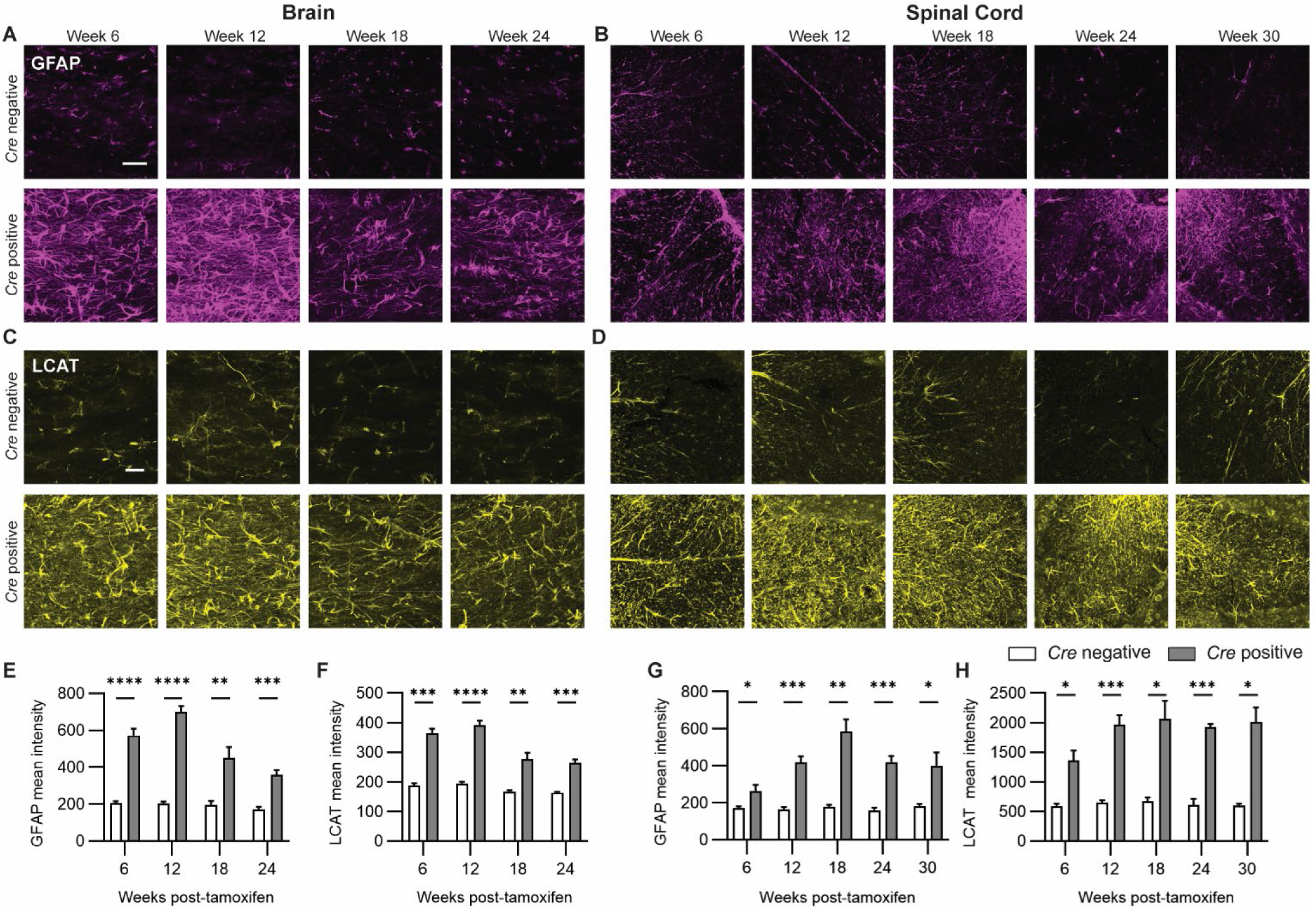
Astrocyte activation parallels LCAT expression. (**A** and **C**) Representative immunofluorescence images in the brain medial corpus callosum. Scale bars indicate 50 µm. Brain sections were stained with antibodies for GFAP (magenta) and LCAT (yellow). (**E** and **F**) Mean intensities in the corpus callosum were obtained using the Fiji ImageJ software. For GFAP analysis, the number of animals analyzed for each group were the following: *Cre* negative, 6 weeks n = 5, 12 weeks n = 6, 18 weeks n = 6, and 24 weeks n = 5; *Cre* positive, 6 weeks n = 5, 12 weeks n = 6, 18 weeks n = 5, and 24 weeks n = 7. For LCAT analysis, n = 4 animals were analyzed for all groups. (**B** and **D**) Immunofluorescence images in the dorsal region of the lumbar spinal cord. Sections were stained with antibodies for GFAP (magenta) and LCAT (yellow). (**G** and **H**) Mean intensities in the dorsal white matter were obtained using the Fiji ImageJ software. For GFAP analysis, two spinal cord sections were stained and the intensities averaged for each animal. The number of animals analyzed were the following: *Cre* negative, 6 weeks n = 5, 12 weeks n = 5, 18 weeks n = 5, 24 weeks n = 4, and 30 weeks n = 5; *Cre* positive, 6 weeks n = 7, 12 weeks n = 8, 18 weeks n = 7, 24 weeks n = 8, and 30 weeks n = 5. For LCAT analysis, n = 3-4 animals were analyzed for all groups. Statistical analysis was performed with multiple t tests comparing *Cre* negative and *Cre* positive at each time point using a Holm–Šídák correction for multiple comparisons (*P < 0.05; **P < 0.01; ***P < 0.001; ****P < 0.0001). For all images, scale bar represents 50 µm.

In the CNS, LCAT is secreted by astrocytes and has been linked to lipid metabolism in neurodegenerative diseases;^25, 26^ however, the role of LCAT in the CNS during demyelination is not well understood. Using immunofluorescence, we observed that LCAT levels were significantly increased in the brain (Figure 6C and 6F) and spinal cord (Figure 6D and 6H) with demyelination. Co-localization analysis of LCAT in astrocytes showed that 76% of brain LCAT and 87% of spinal cord LCAT was expressed in astrocytes during demyelination, confirming that LCAT was primarily an astrocytic protein (Figure 7). The marked overexpression of LCAT in response to demyelination suggests that the LCAT likely also contributes to the observed formation of cholesterol esters. Collectively, our results suggest that multiple biochemical pathways may be responsible for generating cholesterol esters during demyelination and future studies are required to discern how ACAT1 and LCAT each contribute to myelin lipid debris processing.

**Figure 7.**
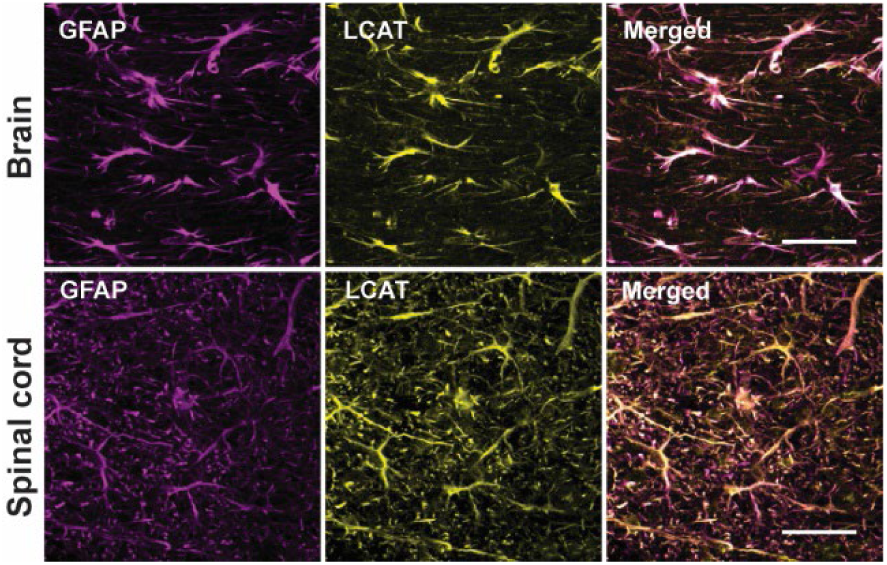
LCAT is expressed in astrocytes. LCAT expression strongly co-localized with GFAP expression. (**A** and **B**) Brain and spinal cord co-localization of LCAT in astrocytes. Scale bars indicate 50 µm. Staining was performed using antibodies for GFAP (magenta) and LCAT (yellow). Threshold analysis was carried out to quantify the percentage of LCAT staining that overlapped with GFAP staining. Only *Cre* positive mice used for this analysis. Mice numbers: Brain 6 weeks n = 4, Spinal cord 18 weeks n = 4.

### OPC and oligodendrocytes dynamics during demyelination and remyelination

By effectively clearing myelin debris, microglia and monocyte-derived macrophages can enhance OPC proliferation and differentiation and contribute to remyelination.^40^ Therefore, for a comprehensive analysis of the cellular response to fluctuations in cholesterol ester levels in the CNS, we quantified both OPCs (PDGFRα) and oligodendrocytes (ASPA or GST-pi) during demyelination and remyelination. We observed significant increases in the OPC and oligodendrocyte populations in the brain with demyelination, which declined gradually during remyelination in the brain (Figure S5-S6). This suggests successful recruitment and proliferation of OPCs increased simultaneously with demyelination to facilitate effective oligodendrocyte differentiation and remyelination. In the spinal cord, the OPC population increased more dramatically (6-fold versus 2-fold in the brain) peaking at week 18 and then gradually declining through week 30, but the increase in OPCs did not lead to robust remyelination (Figures S5). Consistent with these findings, the number of oligodendrocytes in the spinal cord did not significantly increase at any timepoint (Figure S6), although there was a high amount of variability amongst the samples that may be related to the extensive demyelination observed.

### Treatment with Sob-AM2 (remyelinating drug) reduces accumulated cholesterol esters in the brain

Sob-AM2 is a thyroid hormone agonist prodrug with excellent blood–brain barrier permeability,^41^ which has emerged as a promising therapeutic agent for promoting remyelination within the CNS. Sob-AM2 has previously been shown to significantly enhance remyelination in the brain of the same mouse model used in this study (*Plp1*-iCKO-*Myrf*).^28^ Building on this, our recent study examined the lipidomic impact of Sob-AM2 treatment in both brain and spinal cord tissues. Notably, Sob-AM2 effectively restored dysregulated lipid levels in the brain, normalizing the elevated levels of key phospholipids, including phosphatidylethanolamine (PE), phosphatidylserine (PS), and phosphatidylinositol (PI).^30^

Here we analyzed the effects of Sob-AM2 treatment on cholesterol ester during demyelination and remyelination. The *Plp1*-iCKO-*Myrf* mice were administered chow containing Sob-AM2 (84 μg/kg daily dose) starting at 2 weeks post-tamoxifen with continuous ad lib feeding until the end of the experiment. Mice were euthanized at week 12 and week 18 to collect the brain and spinal cord tissues for cholesterol ester analysis. In the brain, treating with Sob-AM2 significantly reduced the accumulation of total cholesterol esters at peak demyelination (week 12) (Figure 8), effectively lowering levels across all cholesterol ester species—including saturated, monounsaturated, and polyunsaturated forms (Figure S7). In contrast, treating with Sob-AM2 had a small effect on spinal cord cholesterol ester levels at week 12, but this effect disappeared by week 18 (Figure 8 and S8) suggesting that the action of Sob-AM2 is not sufficient to overcome the extensive demyelination and damage in the spinal cord of *Plp1*-iCKO-*Myrf* mice. However, this is likely a model-specific effect and not a tissue-specific effect as Sob-AM2 has shown therapeutic benefit in the spinal cord in an inflammatory model of demyelination (experimental autoimmune encephalomyelitis).^42^

**Figure 8.**
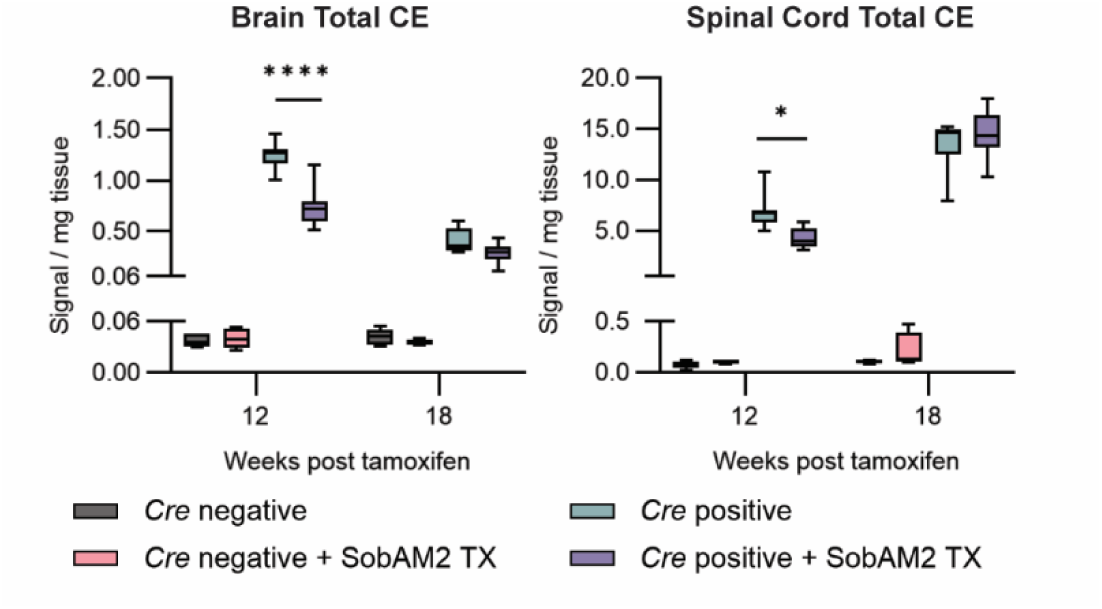
Sob-AM2 treatment limits accumulation of total cholesterol esters levels in the brain. Sob-AM2 reduces the accumulation of cholesterol esters at week 12 in the brain and spinal cord. All individual cholesterol ester species were measured by mass spectrometry and the values were summed for this figure. Statistical analysis was performed using a 3-way ANOVA with a Holm–Šídák test for multiple comparisons (*P < 0.05; ****P < 0.0001). The sample numbers for the brain were the following: *Cre* negative control, 12 weeks n = 5 and 18 weeks n = 6; *Cre* negative SobAM2, 12 weeks n = 6 and 18 weeks n = 4; *Cre* positive control, 12 weeks n = 8 and 18 weeks n =8; *Cre* positive SobAM2, 12 weeks n = 10 and 18 weeks n = 9. The sample numbers for the spinal cord were the following: *Cre* negative control, 12 weeks n = 5 and 18 weeks n = 6; *Cre* negative SobAM2, 12 weeks n = 6 and 18 weeks n = 4; *Cre* positive control, 12 weeks n = 8 and 18 weeks n = 9; *Cre* positive SobAM2, 12 weeks n = 10 and 18 weeks n = 9. Abbreviations: CE, cholesterol esters.

Sob-AM2 has been shown to promote remyelination in the brain^28^ and we now observe reduced cholesterol ester accumulation (Figure 8), which raises several questions about how Sob-AM2 is limiting cholesterol ester buildup during demyelination. Sob-AM2, as a thyroid hormone agonist, is known to act directly on OPCs to promote differentiation and myelination, and one possibility is that the reduction in cholesterol esters could be a secondary byproduct of increased remyelination; faster myelin repair would require more cholesterol, which could happen either by increasing the rate cholesterol ester hydrolysis or bypassing cholesterol ester formation. A second possibility is that Sob-AM2 acts directly on the cholesterol ester formation pathway. Thyroid hormone is known to activate TREM2 and increase microglia phagocytosis,^43^ and direct activation of microglia by Sob-AM2 could lead to increased phagocytosis and flux through the cholesterol ester pathway contributing to the observed improvements in remyelination. Future studies will be needed to determine the mechanism by which Sob-AM2 reduces cholesterol ester metabolism during remyelination.

## Conclusion

This study presents the temporal dynamics of cholesterol esters in the CNS during demyelination and remyelination in the *Plp1*-iCKO-*Myrf* model. The brain and spinal cord both significantly accumulated cholesterol esters during demyelination. In the brain, accumulated cholesterol esters were efficiently cleared with robust remyelination; however, in the spinal cord, demyelination persisted and was accompanied by a chronic accumulation of cholesterol esters highlighting the importance of cholesterol ester clearance for successful remyelination. We also identified that both ACAT1 in microglia and LCAT in astrocytes were overexpressed during demyelination, which suggests that both pathways contributed to the formation of cholesterol esters during demyelination. Formation of lipid droplets were observed in microglia providing further support for a role for microglial ACAT1. Treatment with Sob-AM2, a drug known to promote remyelination in this model, was able to effectively reduce accumulated cholesterol ester levels in the brain further supporting a link between cholesterol ester clearance and remyelination. In conclusion, our study establishes that both cholesterol ester formation and cholesterol ester hydrolysis are critical steps in CNS remyelination and suggests that cholesterol ester metabolism may represent a therapeutic target for promoting robust myelin repair and potentially improving clinical outcomes for people with demyelinating diseases.

## Methods

### Animal Husbandry

Male and female C57BL6/J *Plp1*-iCKO-*Myrf* mice (*Myrf fl/fl*; *Plp1-CreERT*) were generated at the University of Kansas by crossing *Myrf(fl/fl)* mice^44^ with *Plp1-CreERT* mice.^31^ All mice were housed in a climate-controlled environment (24 ± 1 °C) with a constant 12 h light/dark cycle (12 h on, 12 h off) and had unlimited access to food and water. At 8 weeks of age, mice were given daily intraperitoneal injections of 2 mg of tamoxifen (100 μL of 20 mg/mL tamoxifen) in corn oil daily for five consecutive days to induce *Myrf* gene ablation. *Cre* positive mice underwent *Myrf* ablation and demyelination, and *Cre* negative mice served as healthy controls. Mice were euthanized at five timepoints: weeks 6, 12, 18, 24, and 30 post-tamoxifen injections. Mice used for lipid analysis were euthanized by carbon dioxide inhalation and cervical dislocation where brain and spinal cord tissues were collected and immediately, frozen on dry ice, and stored at −80 °C until processing. Mice used for histological analysis were euthanized by intracardiac perfusion with HBSS (Hanks’ Balanced Salt Solution) buffer followed by 4% paraformaldehyde in HBSS.

For the Sob-AM2 drug treatment, control mice received standard chow (Envigo Teklad 2016 diet), and the Sob-AM2 treatment group received the Teklad 2016 diet compounded with 420 μg/kg chow of Sob-AM2 (nominal daily dose of 84 μg/kg body weight) starting 2 weeks after tamoxifen injections. Mice treated with Sob-AM2 were euthanized at 12 and 18 weeks post-tamoxifen by carbon dioxide inhalation and cervical dislocation to collect serum, brain, and spinal cord tissues. All experiments were approved by the Institutional Animal Care and Use Committee at the University of Kansas.

### Lipid Extraction

A modified Bligh–Dyer protocol was used to extract lipids from the brain and spinal cord tissue. Brain tissues were homogenized (either 300 or 50 mg/mL) with ice cold water using a Bead Mill homogenizer (Bead Ruptor Elite, Omni International, USA). Spinal cords were homogenized at a concentration of 65 mg/mL with cold water. Immediately after homogenization, 300 mg/mL brain homogenates were diluted with cold water (3:19) and 950 µLs of the diluted homogenate was used for lipid extractions. For the 50 mg/mL brain homogenates, 1 mL of homogenate was directly used. Spinal cord homogenates were diluted 10-fold and 1 mL was used for the lipid extractions. For all samples, 10 μL of the diluted homogenate was removed and stored at −80 °C for protein quantification using a Bicinchoninic Acid assay. Then the diluted homogenates were added to a mixture of chloroform (with 0.01% butylated hydroxytoluene, BHT): methanol:water (3:2:1) in glass tubes and mixed well with shaking and vortexing. Then the mixture was centrifuged (Sorvall ST 40R Centrifuge, Thermo Fisher Scientific) at 1300 rpm for 10 min and the lower layer was carefully removed and saved in a glass tube. The remaining top layer was further extracted twice with 1.25 mL of chloroform with 0.01% BHT; the lower layers were carefully removed and combined. The combined lower layer was then washed with 300 μL of 1 M KCl followed by 300 μL of water and vacuum-dried completely (Savant SpeedVac SPD130DLX vacuum concentrator, Thermo Fisher Scientific, USA) to obtain the dried lipid extract.

Serum (3 μL) was mixed with 1.2 mL of chloroform: methanol:300 mM ammonium acetate in water (300:665:35) in a glass vial. The dissolved samples were mixed with vortexing and then centrifuged for 5 min at low speed to pellet proteins before mass spectrometry analysis.

### Mass Spectrometric Analysis

Quantification of cholesterol esters was performed by direct infusion triple quadrupole mass spectrometry on a Sciex 4000 QTrap at the Kansas Lipidomic Research Center at Kansas State University. Briefly, dried lipid samples (brain and spinal cord) were dissolved in 1 mL of chloroform. Aliquots were mixed with the internal standard 18:1(d7) cholesterol ester and solvents, and analysis was carried out as previously described.^45^ Cholesterol ester measurements were performed with the use of the acquisition and data processing parameters indicated in Table S2. Cholesterol esters were detected with a precursor ion scan. The fatty acid component was identified based on the number of total acyl carbons and total double bonds, but the individual fatty acids in diacylated lipids, their positions on the glycerol, and double bond positions were not determined. The list of cholesterol ester species analyzed is indicated in Table S2. Data were reported as normalized mass spectral intensity where a value of 1 indicates the intensity of 1 nmol of internal standard (Table S3-S4).

The mass spectrometry analysis was performed in three batches: samples from the week 6, 12, and 18 groups (including control and Sob-AM2 treatment groups) were analyzed at the same time, whereas the week 24 and week 30 groups were analyzed at different times. The brain and spinal cord were analyzed on separate days.

Gas chromatography-mass spectrometry (GC–MS) was used to quantify cholesterol as previously described.^3, 46^ The dried lipid extract was dissolved in 1 mL of chloroform containing 10 nmol of coprosterol as an internal standard and dried under nitrogen. The spiked sample was solubilized with 1 mL of 1:9 3 N potassium hydroxide:methanol and heated at 80 °C for 1 h. After cooling to room temperature, 2 mL of optima-grade water and 0.25 mL of saturated sodium 74 chloride were added. The solution was extracted three times with 2 mL of hexane. All hexane layers were combined, dried under nitrogen, and dissolved in 50 μL of pyridine. Cholesterol derivatization was performed by adding N-trimethylsilyl-N-methyltrifluoroacetamide with trimethylchlorosilane (25 µL) and heating at 50 °C for 1 h. The samples were analyzed on an Agilent 6890 N GC coupled to an Agilent 5975 N quadrupole mass selective detector with electron ionization. The GC contained a VF-5MS capillary column (inert 5 % phenylmethyl column, length: 30 mm, internal diameter: 250 μm, film thickness: 0.25 μm). The carrier gas was helium (1 mL/min). The mass spectrometer was operated in the electron impact mode at 70 eV ionization energy. The MS source temperature was at 230 °C, the front inlet was 250 °C, and the quadrupole temperature was 150 °C. An autosampler (Agilent 7683) was used to the derivatized sample (1 μL) in the splitless mode. The GC had an initial temperature of 150 °C, held for 1 min, ramped 30 °C/min to 300 °C, and then ramped 3 °C/min to a final temperature of 315 °C, and held for 4 min with a total run time of 15 min. The data acquisition was in scan mode (m/z 50 to 650), and the data were processed with Agilent Chemstation software.

### Black-Gold II Staining

Free floating sections (40 μm) of coronal brain sections (bregma 0.020 - 1.045 mm) and lumbar spinal cord were obtained using a Vibratome (Leica VT 1200S). Sections were mounted on positively charged slides and dried overnight at room temperature. Black-Gold staining was performed according to the manufacturer’s protocol with the following modifications. Mounted sections were rehydrated in deionized water for 2 min. Then the slides were transferred into a 0.3% Black-Gold II (Histo-Chem) solution for 3– 5 h at room temperature until desired myelin impregnation was observed. Then the slides were rinsed with deionized water and transferred into a 1% sodium thiosulfate solution for 3 min. Slides were again rinsed with deionized water three times. Sections were dehydrated for 30 s in each ethanol solution (50, 75, 85, 95, 100%). Finally, the slides were immersed in xylene for 1–2 min and secured with coverslips using Cytoseal XYL mounting media (Thermo Fisher). Mounted slides were heated on a slide warmer for 2–3 h at 60 °C. Sections were imaged using a slide reader (BioTek Cytation 5) at 4× magnification. For the stained brain tissues, multiple images were taken and stitched together to obtain the whole image.

Black-Gold images were analyzed by threshold analysis to determine a percent myelin staining within the tissue. For the brain tissue staining, all timepoints (week 6,12,18 and 24) were stained together. For the spinal cord tissue staining, each timepoint (weeks 6,12, 18,24 and 30) was stained as a separate batch. Thresholds were set for each batch using *Cre* negative images to encompass the major white matter tracts (brain: 0–50; and spinal cord: 0–100 for week 6, 0‒80 for week 12, 0‒75 for week 18, 0‒90 for week 24 and 0–105 for week 30). Direct comparisons were only performed on sections stained in the same batch. Statistical analysis was performed to compare *Cre* negative and *Cre* positive tissues at each time point using unpaired *t* tests corrected for multiple comparisons using the Holm–Šídák method. Brains were analyzed from *Cre* negative mice and *Cre* positive mice (*n* = 3 at week 6,12,18 and 24). Spinal cords were analyzed from *Cre* negative mice (*n* = 6 at week 6 and 12, *n* = 5 at week 18, 24 and 30) and *Cre* positive mice (*n* = 7 at week 6, *n* = 12 at week 12, *n* = 8 at week 18, *n* = 6 at week 24 and *n* = 5 at week 30).

### Immunofluorescence staining

Free floating sections (40 μm) were obtained from brain (bregma 0.020 - 1.045 mm) or lumbar spinal cord using Vibratome (Leica VT 1200S). Sections were rinsed two times with HBSS/0.01% sucrose. Sections were then permeabilized using 0.05% Triton X-100 in HBSS with 0.01% sucrose and 0.1% saponin (HBSS/Su/Sap) for 30 minutes at room temperature followed by rinsing using HBSS/Su/Sap for 10 minutes. Sections were incubated in blocking buffer (3% normal donkey serum diluted in HBSS/Su/Sap) for 2 hours at room temperature. Then sections were treated with primary antibodies at room temperature for 3.5 hours and further incubated at 4 °C overnight with shaking. Tissue sections were rinsed with HBSS buffer and incubated with secondary antibodies for 3-4 hours at room temperature and incubated with DAPI for another 1 hour. Finally, the sections were rinsed and mounted on slides using ProLong Diamond antifade mounting media (Life Technologies P36961).

Primary antibodies used were specific for GFAP (EMD Millipore AB5541, 1:500), IBA1 (Fujifilm Wako 019-19741, 1:500), PDGFRα (R&D Systems AF1062, 1:500), Aspa/Nur7 (Millipore ABN1698, 1:100), GST-pi (MBL Life Science 312, 1:500), ACAT1 (Proteintech 16215-1-AP, 1:50), LCAT (Proteintech 12243-1-AP, 1:50. Secondary antibodies were from Life Technologies (A31572, A21447, A48272TR, A78952, all at 1:200).

For the lipid droplet staining free floating sections (40 µm) were first permeabilized using 0.05% Triton X-100 in PBS. Then the brain and spinal cord sections were stained using LipidSpot 610 (Biotium, 1:1000) or Lipidspot 488 (Biotium, 1:1000), respectively, for 30 minutes protected from light. Stained sections were rinsed and mounted on slides using PBS and images were acquired on the same day.

Co-staining for both IBA1 and ACAT1 was carried out by directly labeling the primary antibodies using Mix-n-Stain Antibody labeling kits. The ACAT1 antibody (Proteintech 16215-1-AP) was conjugated with Mix-n-Stain CF555 (Biotium 92254, 20-50 µg), and the IBA1 antibody (Fujifilm Wako 019-19741) was conjugated with Mix-n-Stain CF647 (Biotium 92259, 20-50 µg). After the antibody conjugates were made, brain and spinal cord sections were stained at room temperature for 3.5 hours and further incubated at 4 °C overnight with shaking. Sections were rinsed and mounted on slides using ProLong Diamond antifade mounting media.

Immunofluorescence-stained sections were imaged using a Nikon 2037 CSU-W1 spinning disk confocal microscope at the KIDDCR imaging Coret the University of Kansas Medical Center. Brains were imaged at the medial corpus callosum and spinal cords were imaged at the ventral or dorsal medial white matter regions as indicated. For most analyses, the sections were imaged with a ×20/NA 0.8 objective (710 × 744 μm), and Z-stacks were acquired every 4 µm. For the co-localization studies (IBA1-ACAT1, GFAP-LCAT, and lipid droplet-IBA1-GFAP), sections were imaged with a 4×SoRa objective (177 x 186 μm) and Z-stacks were obtained every 0.8 μm. For analysis, the Z-stacks were compressed into a single maximum intensity projection image.

A region of interest was drawn around the medial corpus callosum or the spinal cord medial white matter for quantification, and the full images were cropped and enlarged for clarity in Figures 3, 5, and 6 as indicated in Figure S9. PDGFRα-labeled cells within the region of interest were manually counted and ASPA and GST-pi-labeled cells were counted using an automated method using Fiji ImageJ software. For the analysis of IBA1, GFAP, ACAT1, LCAT, and lipid droplets, mean intensity of staining in the region of interest was calculated. Colocalization analysis was performed by thresholding the images to generate a binary image of the area of expression for each marker. Then the area of overlap between the two markers was quantified and divided by the area of the cell type of interest (IBA1 for microglia or GFAP for astrocytes) to obtain a % colocalization.

### Electron Microscopy

For the electron microscopy analysis, brain and spinal cord samples from week 12 and week 24 that were initially perfused with 4% paraformaldehyde in HBSS. The embedding of the brain and spinal cord sections was performed by the University of Kansas Microscopy and Analytical Imaging Core Laboratory. Blocks for plastic embedding were dissected from the corpus callosum in the brain (bregma -0.955 to 0.020 mm) and 500 μm cross sections were dissected from the lumbar region of the spinal cord. Seven days prior to embedding, brain and spinal cord samples were immersed individually in microtubes with Karnovsky’s fixative at 4 °C, which contained 2.5% glutaraldehyde/ 2% formaldehyde^47^ diluted in 0.2 M sodium cacodylate buffer (pH 7.4) + 0.01 M sucrose (SCB/Su). Next, tissue samples were rinsed once with SCB/Su for 5 min to continue the procedure using a laboratory microwave (PELCO BioWave® Pro 36500; Ped Pella, Inc.). First, samples were post-fixed in a modified Karnovsky’s fixative (2.5% glutaraldehyde/ 2% formaldehyde^47^) diluted in SCB/Su, rinsed with SCB/Su, and stained with 2% osmium tetroxide diluted SCB/Su. Then, samples were rinsed with 50% followed by 70% ethanol diluted in carbon dioxide-free deionized water. Next, tissues were en bloc stained with 4% uranyl acetate + 0.4% lead citrate^48^ diluted in 70% ethanol carbon dioxide-free water. Next, samples were dehydrated with 95% ethanol and en bloc stained with 1% alcoholic phosphotungstic acid^49^ diluted in 100% dry ethanol + 5 drops of 95% ethanol, continuing dehydration with 100% dry ethanol, 1:1 100% dry ethanol: 100% dry acetone, followed by Embed 812 resin infiltration steps (3:1 100% dry acetone: Embed 812 resin; 1:1 100% dry acetone: Embed 812 resin; 1:3 100% dry acetone: Embed 812 resin, 100% Embed 812 resin and a second change of 100% Embed 812 resin). Samples remained immersed in Embed 812 resin overnight at RT inside the fume hood. Next day, each sample was moved individually on a flat mold filled with embedding resin (Embed 812 resin + 2,4,6-tri(dimethylaminomethyl)phenol and placed inside an oven at 65 °C for 60-72 hours.

The embedded brain and spinal cords were sectioned and imaged by the Oregon Health Science University (OHSU) Multiscale Microscopy Core (MMC). For the examination of corpus callosum, semi-thick sections (500 nm) were collected and stained with toluidine blue. Sections were scanned using a uSCOPE MXII (Microscope International, LLC.), equipped with a 20x/0.4 NA 160 mm conjugated objective lens. Images were acquired using uScope Navigator 4.6. Image stitching was performed using Fiji/ImageJ 1.54p. Sample blocks were further trimmed, ultrathin sections (100 nm) were collected and counterstained with 2% aqueous uranyl acetate and Reynold’s lead citrate. Sections were examined with a FEI Tecnai T12 transmission electron microscope, operated at 80 kV, and digital images were acquired with an AMT Nanosprint12 4k x 3k camera. Brains were imaged at a direct magnification of 6800x and spinal cords were images at a direct magnification of 2900x.

*Cre* negative and *Cre* positive brain and spinal cord samples at weeks 12 and 24 were analyzed with n = 3-4 mice for all groups. The percentage of myelin area was determined for 3 images per mouse using threshold analysis. The threshold was set from 0-125, which encompassed the myelin membranes for most of the images. In select images (∼10%) that were either darker or lighter overall, the threshold was adjusted to 0-115 or 0-135, respectively, to ensure that the thresholded area represented the myelin regions. All adjustments were made in a blinded manner. In addition, g-ratio analysis was performed by measuring the axon circumference (inside the myelin sheath) and the myelin circumference (outside the myelin sheath). The g-ratio was calculated by dividing the inner circumference by the outer circumference. Three distinct images were analyzed for each mouse and 30-50 axons were quantified per image (a minimum of 100 axons per mouse).

### Statistical Analysis

All comparisons of *Cre* negative (healthy) and *Cre* positive (demyelination) across timepoints were analyzed with multiple t tests using a Holm–Šídák correction for multiple comparisons (^∗^P < 0.05; ^∗∗^P < 0.01; ^∗∗∗^P < 0.001; ^∗∗∗∗^P < 0.0001). For the cholesterol ester measurements that considered Sob-AM2 treatment, statistical analysis was performed with a three-way ANOVA comparing across timepoints, drug treatment, and healthy/demyelination with the Holm–Šídák test for multiple comparisons (^∗^P < 0.05; ^∗∗^P < 0.01; ^∗∗∗^P < 0.001; ^∗∗∗∗^P < 0.0001).

## Abbreviations

ACAT1: acetyl-CoA-acyltransferase_1
CE: cholesterol ester
CNS: central nervous system
iCKO: induced conditional knockout
LCAT: lecithin-cholesterol acyltransferase, LCAT
*Myrf*: myelin regulatory factor
OPC: oligodendrocyte precursor cell

## Acknowledgements

Confocal microscopy was performed at the Integrative Imaging Core Laboratory at the University of Kansas Medical Center (Kansas City, KS) and was supported by NIH S10 OD032207. Samples for electron microscopy were prepared by the Microscopy and Analytical Imaging Core at the University of Kansas (Lawrence, KS). Electron microscopy was performed by Rodolfo Alvarado at the Multiscale Microscopy Core at Oregon Health & Science University (OHSU, Portland, OR), a member of the OHSU Research Cores and Shared Resources, RRID: SCR_009969.

## Author Contributions

Samples were prepared for mass spectrometry analysis by NDSM, RB and DD. Animal work was performed by JMW, RB, and HK. Microscopy and analysis were performed by NDSM. Electron microscopy analysis was performed by JMP, AG, and MDH. NDSM and MDH conceived the project, designed the experiments, analyzed the data, and wrote the manuscript.

## Supplementary Information

Graphs of the total CE levels for serum; graphs of all individual CE species for the brain, spinal cord, brain treated with Sob-AM2, and spinal cord treated with Sob-AM2; representative images of the spinal cord Black-Gold and lipid droplet staining, representative images and quantified analysis of the number of OPCs and oligodendrocytes in brain and spinal cord; schematic of how the regions of interest in the fluorescence images were selected.

## For Table of Contents Only

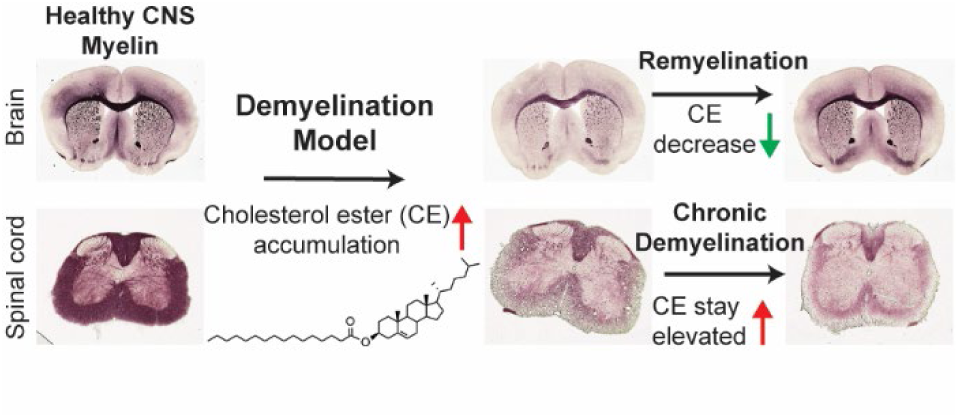

