## Supplemental Figures for "Temporal dynamics of CNS cholesterol esters correlate with demyelination and remyelination" (from another version of this preprint: version 1, 2025-09-11)

**Table S1. Cholesterol ester accumulation represents a major percentage of the total brain cholesterol.**

Tissue samples were analyzed at the time point of maximum cholesterol ester accumulation. Brain tissue was analyzed from 12 weeks post-tamoxifen and spinal cord was analyzed from 18 weeks post-tamoxifen. Data are shown as mean ± standard error of mean (SEM).

| **Tissue** | **Genotype** | **n** | **Cholesterol (nmol/ mg tissue)** | **Cholesterol ester (nmol/ mg tissue)** | **% Cholesterol ester** |
|---|---|---|---|---|---|
| Brain | *Cre* neg (healthy) | 5 | 10.3 ± 0.9 | 0.04 ± 0.01 | 0.4 ± 0.1 |
| Brain | *Cre* pos (demyelination) | 8 | 8.5 ± 0.6 | 1.2 ± 0.1 | 13.0 ± 0.9 |
| Spinal cord | *Cre* neg (healthy) | 6 | 77 ± 3 | 0.10 ± 0.01 | 0.14 ± 0.01 |
| Spinal cord | *Cre* pos (demyelination) | 9 | 43 ± 2 | 13.5 ± 0.8 | 24 ± 1 |

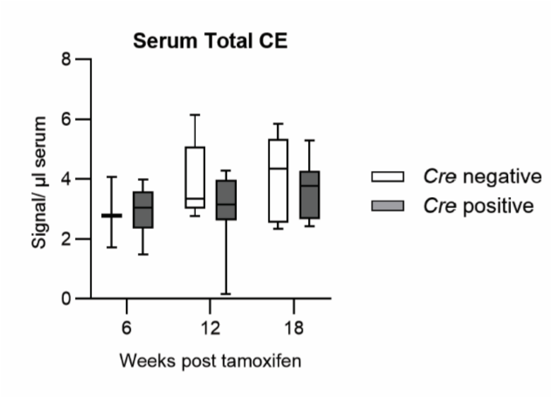


**Figure S1. Total cholesterol ester levels in serum with demyelination and remyelination.**

Total cholesterol ester levels did not change with demyelination and remyelination in the serum. Sample numbers are: *Cre* negative, 6 weeks n = 3, 12 weeks n = 5, and 18 weeks n = 6; *Cre* positive, 6 weeks n = 8, 12 weeks n = 8, and 18 weeks n = 8. Statistical analysis was performed with multiple t tests comparing *Cre* negative and *Cre* positive at each time point using a Holm–Šídák correction for multiple comparisons.

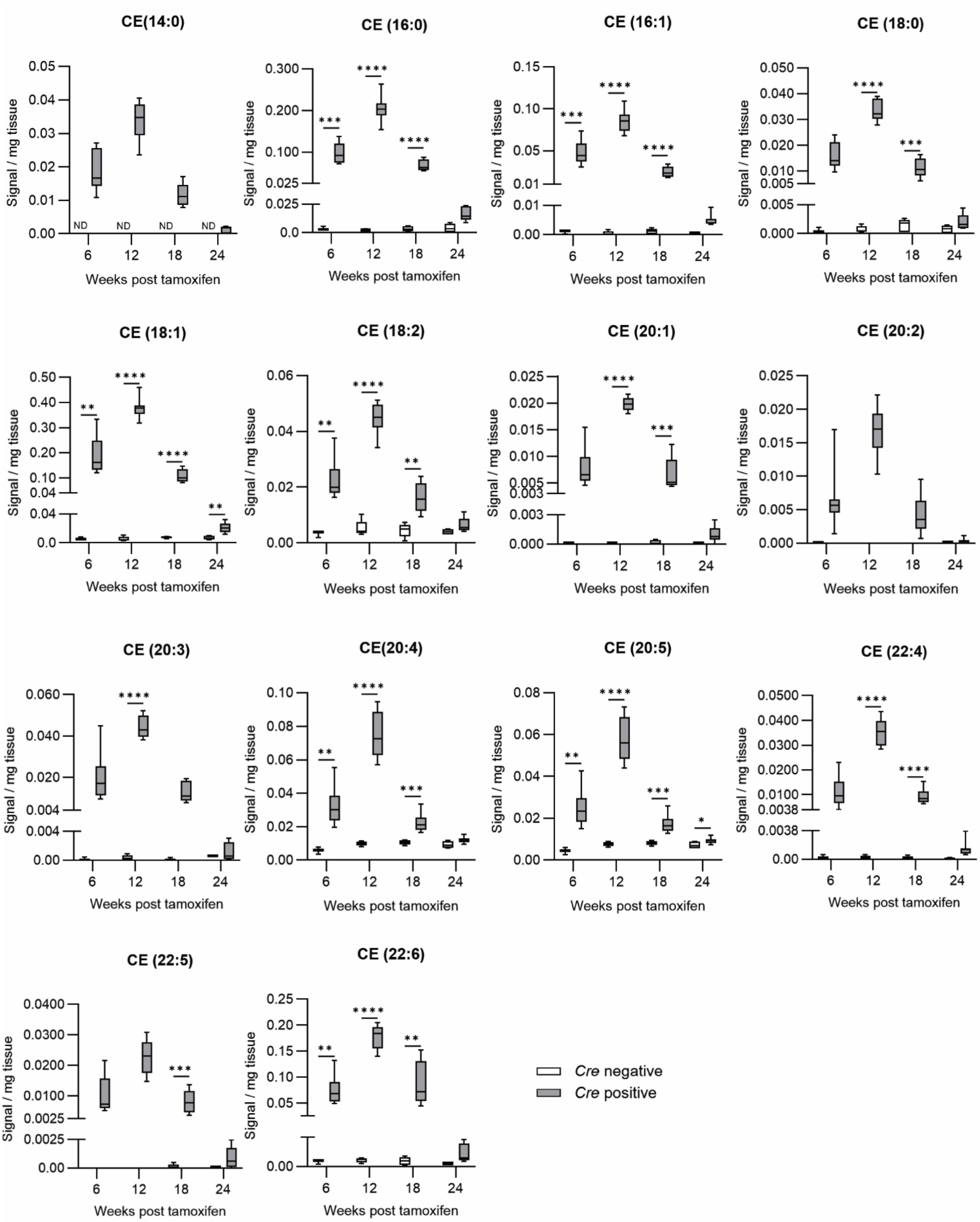

CE(14:0)
Signal / mg tissue
ND
ND
ND
ND
Weeks post tamoxifen
CE (16:0)
Signal / mg tissue
Weeks post tamoxifen
CE (16:1)
Signal / mg tissue
Weeks post tamoxifen
CE (18:0)
Signal / mg tissue
Weeks post tamoxifen
CE (18:1)
Signal / mg tissue
Weeks post tamoxifen
CE (18:2)
Signal / mg tissue
Weeks post tamoxifen
CE (20:1)
Signal / mg tissue
Weeks post tamoxifen
CE (20:2)
Signal / mg tissue
Weeks post tamoxifen
CE (20:3)
Signal / mg tissue
Weeks post tamoxifen
CE(20:4)
Signal / mg tissue
Weeks post tamoxifen
CE (20:5)
Signal / mg tissue
Weeks post tamoxifen
CE (22:4)
Signal / mg tissue
Weeks post tamoxifen
CE (22:5)
Signal / mg tissue
Weeks post tamoxifen
CE (22:6)
Signal / mg tissue
Weeks post tamoxifen
Cre negative
Cre positive

**Figure S2. All cholesterol ester species in the brain increase during demyelination and decreases during remyelination.**

The most abundant cholesterol ester species were quantified by mass spectrometry. The data are plotted as box and whisker plots with the bars representing the minimum and maximum values. The sample numbers are the following: *Cre* negative, 6 weeks n = 3, 12 weeks n = 5, 18 weeks n = 6, and 24 weeks n = 4; *Cre* positive, 6 weeks n = 8, 12 weeks n = 8, 18 weeks n = 8, and 24 weeks n = 7. Statistical analysis was performed with multiple t tests comparing *Cre* negative and *Cre* positive at each time point using a Holm–Šídák correction for multiple comparisons (*$P < 0.05$; **$P < 0.01$; ***$P < 0.001$; ****$P < 0.0001$).

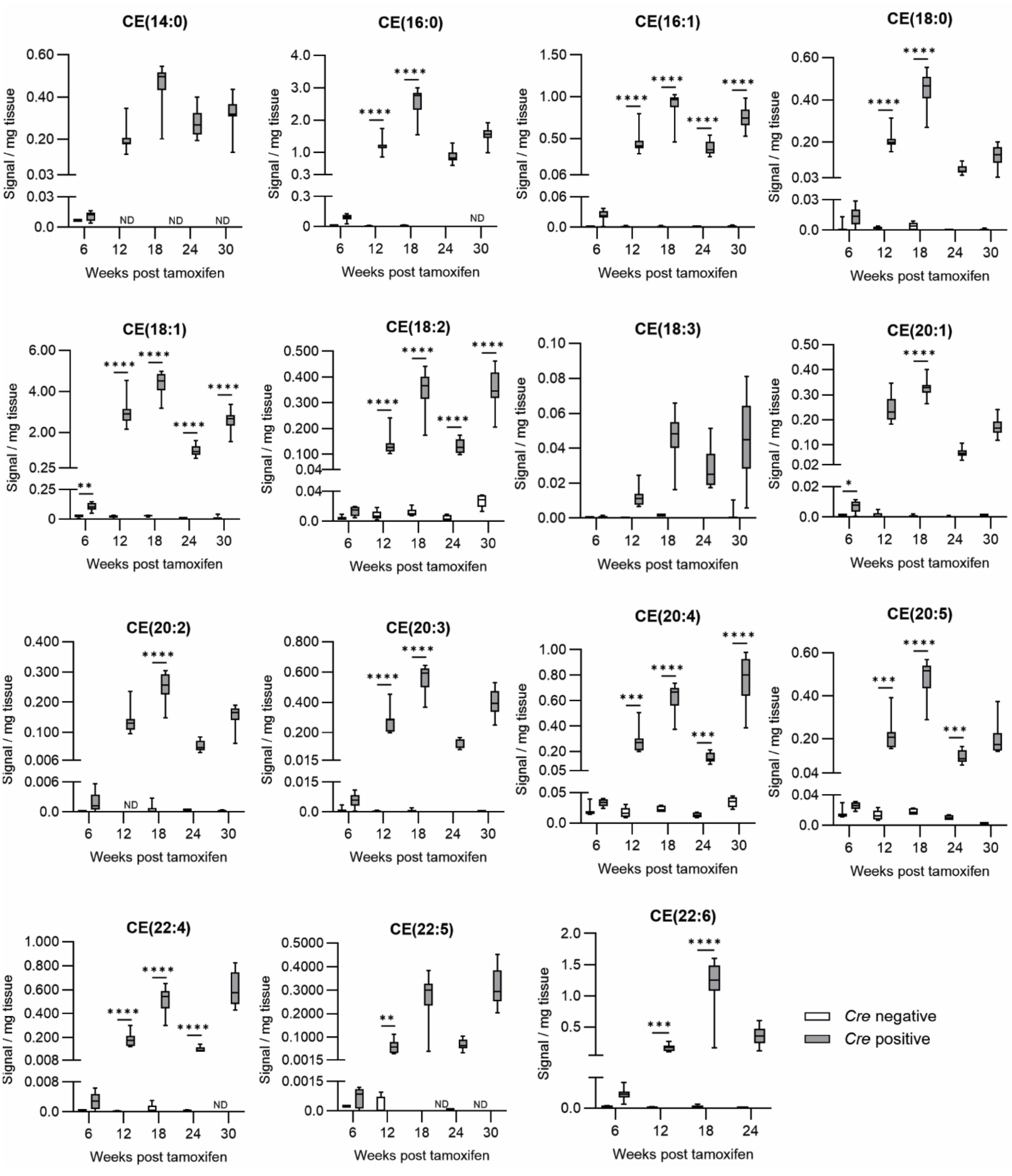


**Figure S3. Cholesterol ester species in the spinal cord remain elevated at all time points.**

The most abundant cholesterol ester species were quantified by mass spectrometry. The data are plotted as box and whisker plots with the bars representing the minimum and maximum values. The sample numbers are the following: *Cre* negative, 6 weeks $n = 3$, 12 weeks $n = 5$, 18 weeks $n = 6$, 24 weeks $n = 4$, and 30 weeks n=7; *Cre* positive, 6 weeks $n = 8$, 12 weeks $n = 8$, 18 weeks $n = 9$, 24 weeks $n = 7$, and 30 weeks n = 7. Statistical analysis was performed with multiple *t* tests comparing *Cre* negative and *Cre* positive at each time point using a Holm–Šídák correction for multiple comparisons (**$P < 0.01$; ***$P < 0.001$; ****$P < 0.0001$).

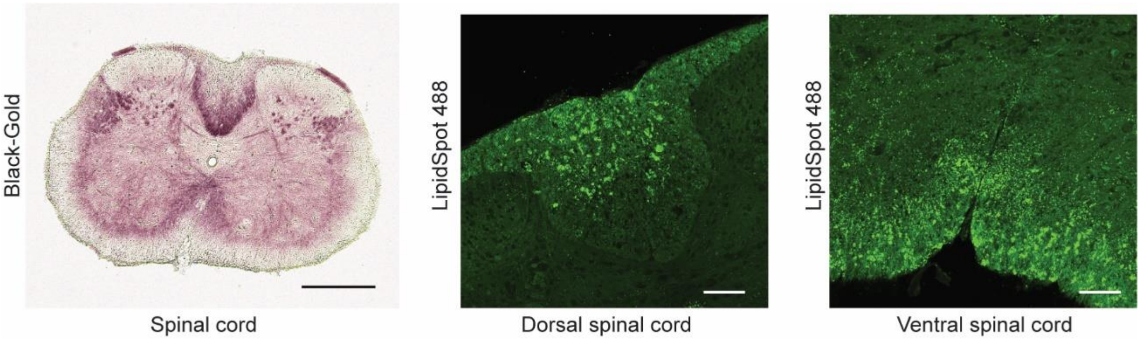


**Figure S4. Lipid droplet accumulation correlated with myelin damage in the outer white matter of the spinal cord.**

Representative spinal cord images from *Cre* positive (demyelinated) 18 weeks post-tamoxifen mice. The most reduced regions of Black-Gold staining are the outer white matter of the dorsal, ventral, and lateral white matter tracts (left), which overlays with the regions of greatest lipid droplet accumulation (right) as measured by staining with LipidSpot 488. The scale bar for the Black-Gold image represents 0.5 mm and the scale bars for the LipidSpot images represent 0.1 mm.

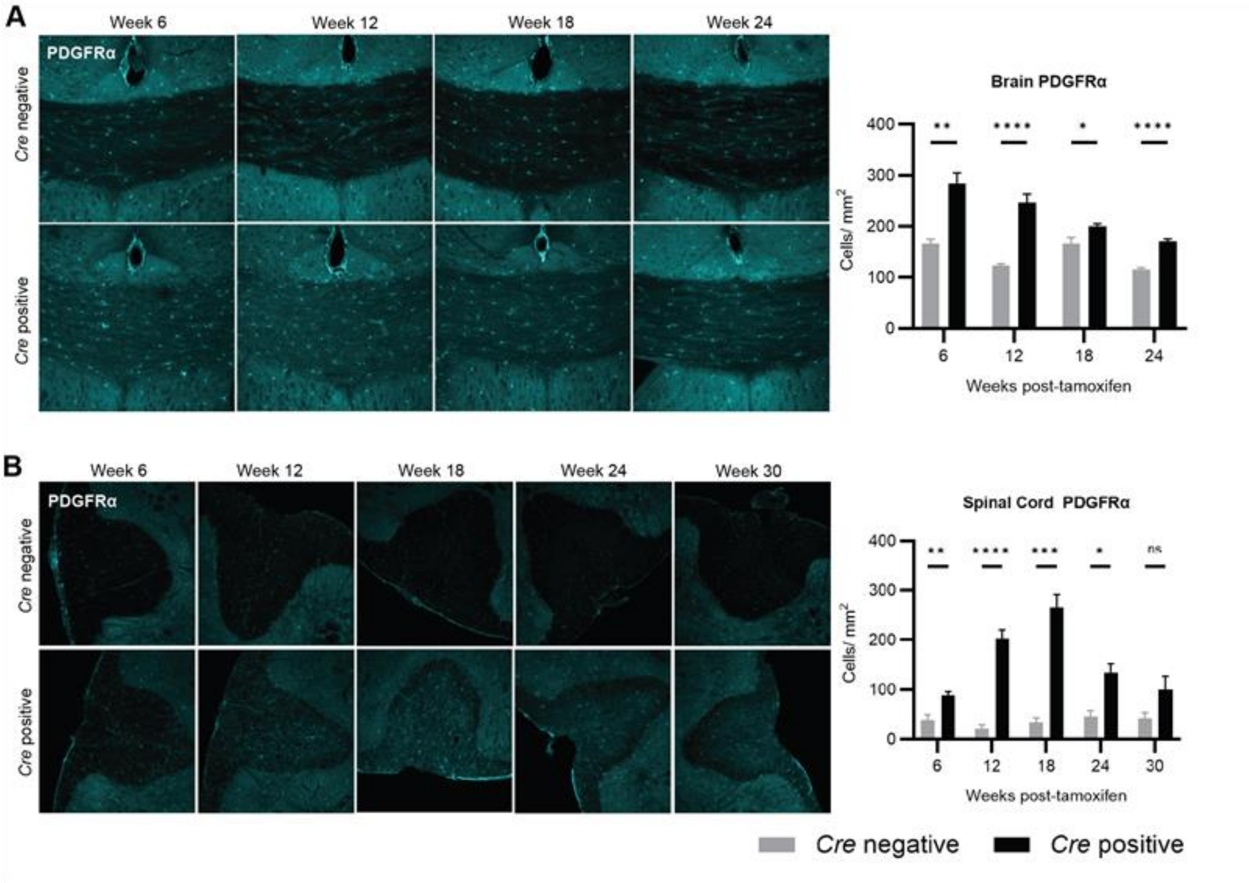


**Figure S5. Oligodendrocyte precursor cell (OPC) response with demyelination and remyelination.**

(A and B) Representative immunofluorescence images in the brain medial corpus callosum and the lumbar spinal cord dorsal white matter. Sections were stained for OPCs with antibodies for PDGFRα (cyan). PDGFRα+ cells were quantified using the cell counter in Fiji ImageJ software. Animal numbers used for this analysis were the following: brain *Cre* negative, 6 weeks n = 5, 12 weeks n = 6, 18 weeks n = 5, and 24 weeks n = 5; brain *Cre* positive, 6 weeks n = 5, 12 weeks n = 6, 18 weeks n = 6, and 24 weeks n = 7; spinal cord *Cre* negative, 6 weeks n = 5, 12 weeks n = 6, 18 weeks n = 5, 24 weeks n = 4, and 30 weeks n = 5; spinal cord *Cre* positive, 6 weeks n = 7, 12 weeks n = 9, 18 weeks n = 8, 24 weeks n = 7, and 30 weeks n = 5. Statistical analysis was performed with multiple t tests comparing *Cre* negative and *Cre* positive at each time point using a Holm–Šídák correction for multiple comparisons (*$P < 0.05$; **$P < 0.01$; ***$P < 0.001$; ****$P < 0.0001$).

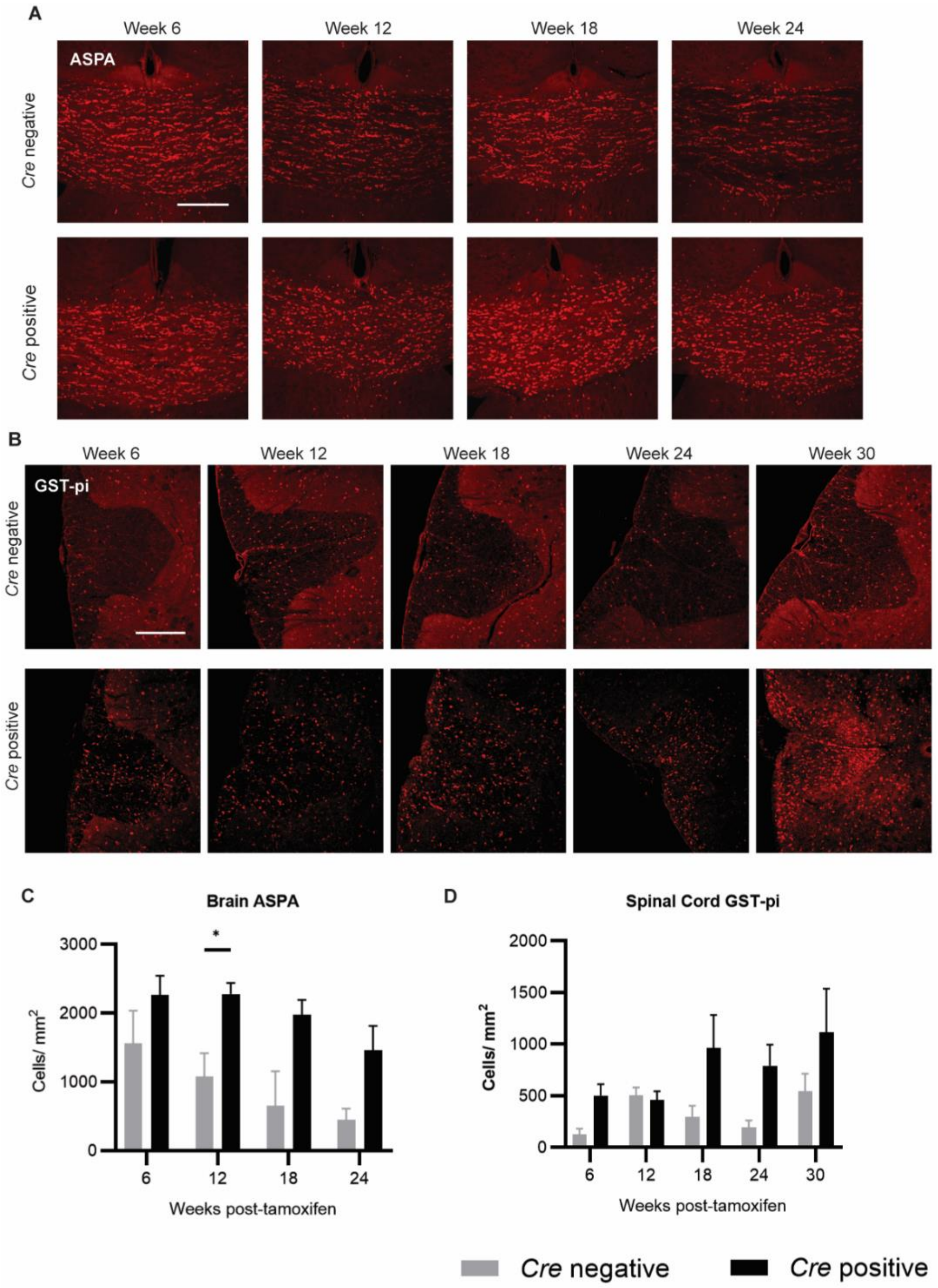


**Figure S6. Oligodendrocyte cell response with demyelination and remyelination.**

(A and C) Representative immunofluorescence images and the quantitative oligodendrocyte cell counts in the brain medial corpus callosum. Brain sections were stained for oligodendrocytes with an antibody for ASPA (red). ASPA+ cells were quantified using the automated cell counting in Fiji ImageJ software. Animal numbers used for this analysis were the following: *Cre* negative, 6 weeks n = 5, 12 weeks n = 6, 18 weeks n = 5, and 24 weeks n = 5; *Cre* positive, 6 weeks n = 5, 12 weeks n = 6, 18 weeks n = 6, and 24 weeks n = 6. (B and D) Representative immunofluorescence images and the quantitative oligodendrocyte cell counts in the lumbar spinal cord dorsal white matter. Spinal cord sections were stained for oligodendrocytes with an antibody for GST-pi (red). GST-pi+ cells were quantified using the automated cell counting in Fiji ImageJ software. Animal numbers used for this analysis were the following: *Cre* negative, 6 weeks n = 5, 12 weeks n = 5, 18 weeks n = 4, 24 weeks n = 5, and 30 weeks n = 3; *Cre* positive, 6 weeks n = 7, 12 weeks n = 9, 18 weeks n = 8, 24 weeks n = 6, and 30 weeks n = 4. Statistical analysis was performed with multiple t tests comparing *Cre* negative and *Cre* positive at each time point using a Holm–Šídák correction for multiple comparisons (*$P < 0.05$)

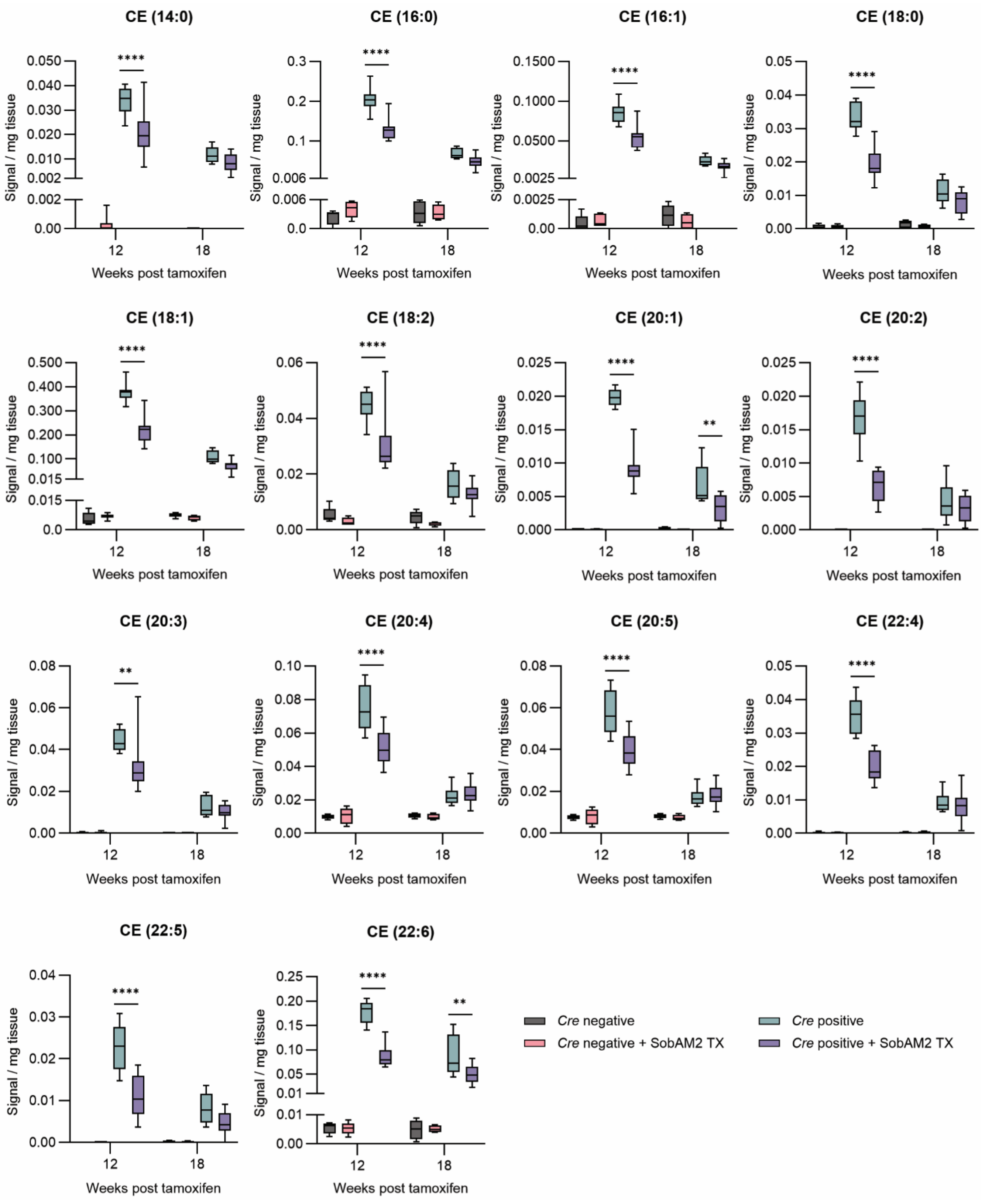


**Figure S7. Individual cholesterol ester species levels in brain tissue at 12 and 18 weeks post tamoxifen with Sob-AM2 treatment (TX).**

The most abundant cholesterol ester species were quantified by mass spectrometry. The data are plotted as box and whisker plots with the bars representing the minimum and maximum values. Statistical analysis was performed with a three-way ANOVA comparing across time, treatment, and genotype with the Holm–Šídák multiple comparisons test (*$P < 0.05$; ***$P < 0.001$; ****$P < 0.0001$). The sample numbers are the following: *Cre* negative control, 12 weeks n = 5 and 18 weeks n = 6; *Cre* negative SobAM2, 12 weeks n = 6 and 18 weeks n = 4; *Cre* positive control, 12 weeks n = 8 and 18 weeks n = 8; *Cre* positive SobAM2, 12 weeks n = 10 and 18 weeks n = 9.

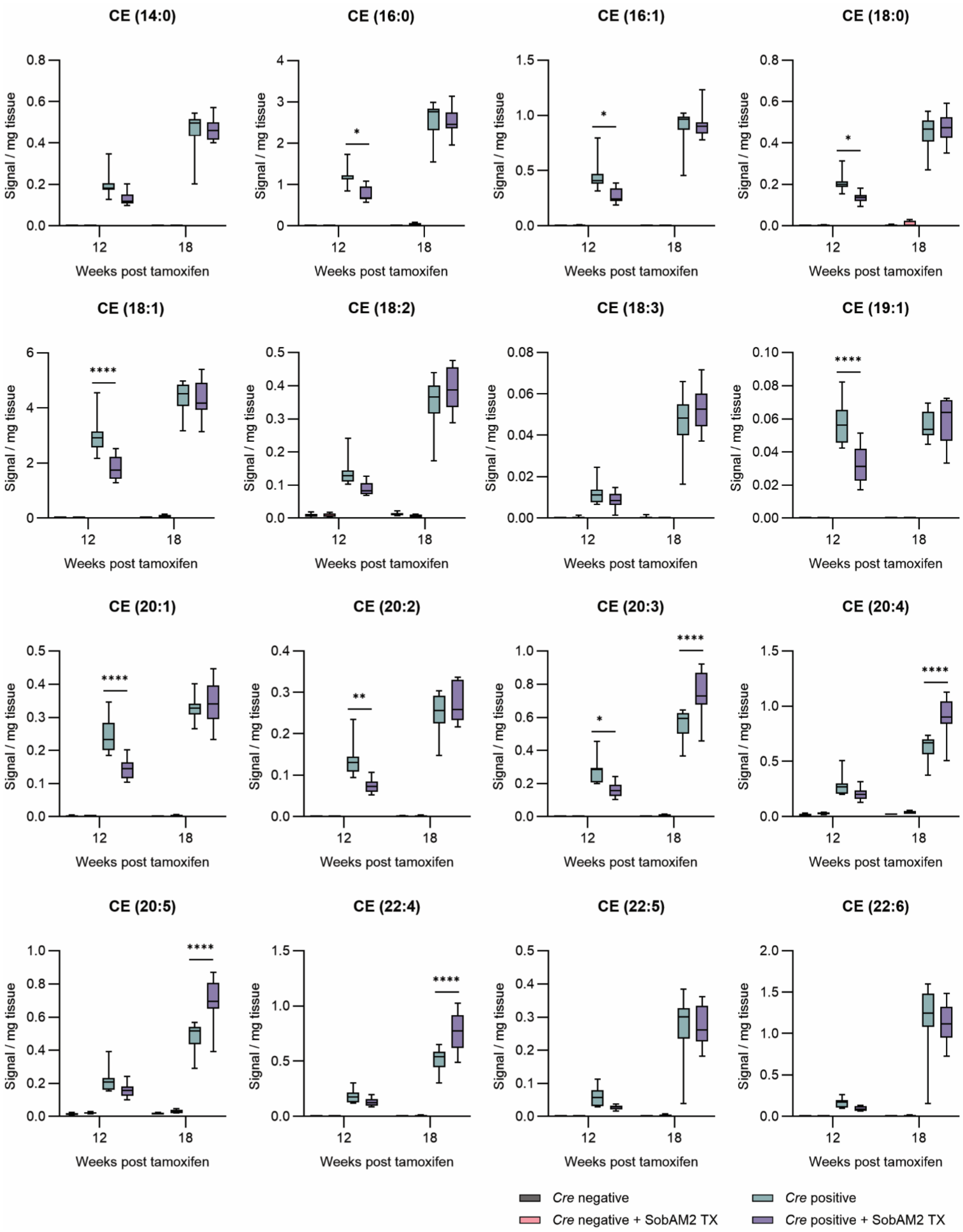
CE (14:0)
CE (16:0)
CE (16:1)
CE (18:0)
CE (18:1)
CE (18:2)
CE (18:3)
CE (19:1)
CE (20:1)
CE (20:2)
CE (20:3)
CE (20:4)
CE (20:5)
CE (22:4)
CE (22:5)
CE (22:6)
Signal / mg tissue
Weeks post tamoxifen
12
18
*
**
****
Cre negative
Cre negative + SobAM2 TX
Cre positive
Cre positive + SobAM2 TX

**Figure S8. Individual cholesterol ester species levels in the spinal cord tissue at 12 and 18 weeks post tamoxifen with Sob-AM2 treatment (TX).**

The most abundant cholesterol ester species were quantified by mass spectrometry. The data are plotted as box and whisker plots with the bars representing the minimum and maximum values. Statistical analysis was performed with a three-way ANOVA comparing across time, treatment, and genotype with the Holm–Šídák multiple comparisons test (*$P < 0.05$; ***$P < 0.001$; ****$P < 0.0001$). The sample numbers are the following: *Cre* negative control, 12 weeks n = 5 and 18 weeks n = 6; *Cre* negative SobAM2, 12 weeks n = 6 and 18 weeks n = 4; *Cre* positive control, 12 weeks n = 8 and 18 weeks n = 9; *Cre* positive SobAM2, 12 weeks n = 10 and 18 weeks n = 9.

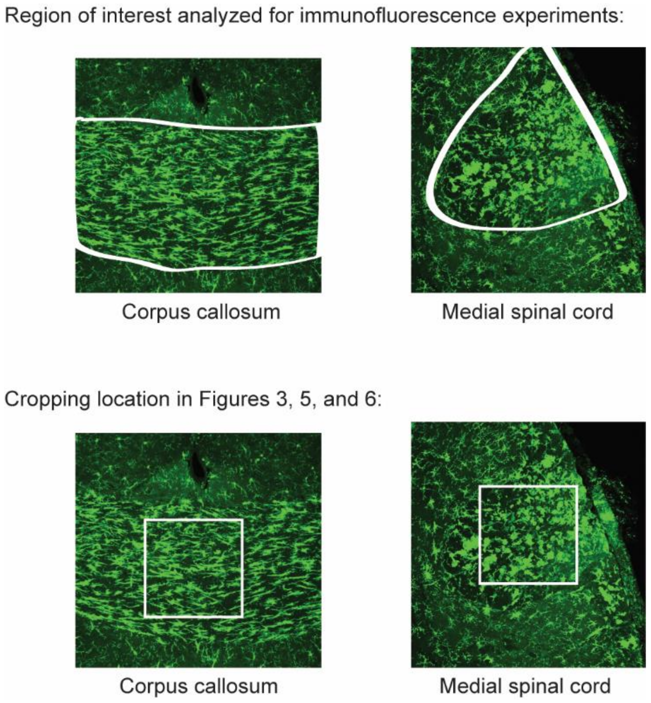


**Figure S9: Region of interest (ROI) analyzed for immunofluorescence experiments**

(Top) The regions of interest analyzed in the medial corpus callosum and medial spinal cord for immunofluorescence experiments are indicated by the white outlines. (Bottom) For Figures 3, 5, and 6, the images were cropped for clarity and the approximate crop locations are shown above (white squares).
